# Conceptual representations in the default, control and attention networks are task-dependent and cross-modal

**DOI:** 10.1101/2023.04.15.536954

**Authors:** Philipp Kuhnke, Markus Kiefer, Gesa Hartwigsen

## Abstract

Conceptual knowledge is central to human cognition. Neuroimaging studies suggest that conceptual processing involves modality-specific and multimodal brain regions in a task-dependent fashion. However, it remains unclear (1) to what extent conceptual feature representations are also modulated by the task, (2) whether conceptual representations in multimodal regions are indeed cross-modal, and (3) how the conceptual system relates to the large-scale functional brain networks. To address these issues, we conducted multivariate pattern analyses on fMRI data. 40 participants performed three tasks—lexical decision, sound judgment, and action judgment—on written words. We found that (1) conceptual feature representations are strongly modulated by the task, (2) conceptual representations in several multimodal regions are cross-modal, and (3) conceptual feature retrieval involves the default, frontoparietal control, and dorsal attention networks. Conceptual representations in these large-scale networks are task-dependent and cross-modal. Our findings support theories that assume conceptual processing to rely on a flexible, multi-level architecture.

## 1. Introduction

Conceptual knowledge is crucial for many cognitive abilities, such as word comprehension and object recognition (Lambon Ralph, 2014; van Elk et al., 2014). Previous neuroimaging studies indicate that conceptual processing involves both modality-specific perceptual-motor regions and cross-modal convergence zones (for a meta-analysis, see Kuhnke et al., 2023; for reviews, see Binder & Desai, 2011; Borghesani & Piazza, 2017; Kiefer & Pulvermüller, 2012; Lambon Ralph et al., 2016).

Modality-specific regions represent perceptual-motor features of concepts. While a common terminology is currently lacking in the field, we refer to “perceptual-motor modalities” as the brain’s major input and output channels of perception and action (Kuhnke et al., 2023, 2021). Note that these modalities do not simply correspond to the senses (hence the term “perceptual-motor” and not “sensory”) as they include channels of internal perception (e.g. emotion) as well as motor action (Kiefer and Harpaintner, 2020). We call brain regions “modality-specific” if they represent information related to a single perceptual-motor modality (Barsalou, 2016; Kiefer and Pulvermüller, 2012). For instance, action features are represented in somatomotor regions (Hauk et al., 2004; Tettamanti et al., 2005; Vukovic et al., 2017), while sound features are represented in auditory areas (Bonner and Grossman, 2012; Kiefer et al., 2012, 2008; Trumpp et al., 2013a). These findings support grounded cognition theories, which propose that concepts consist of perceptual-motor features represented in modality-specific perceptual-motor areas (Barsalou, 2008; Pulvermüller, 2012).

In contrast, cross-modal convergence zones integrate modality-specific features into more abstract, cross-modal representations (Binder, 2016; Fernandino et al., 2016a; Kuhnke et al., 2023, 2020b; Tong et al., 2022). We previously proposed a distinction among cross-modal convergence zones between “multimodal” regions which retain modality-specific information, and “amodal” regions which completely abstract away from modality-specific input (Kuhnke et al., 2023, 2022, 2020b). Multimodal regions seem to include the left inferior parietal lobe (IPL) and posterior middle temporal gyrus (pMTG) (Fernandino et al., 2022, 2016a; Kuhnke et al., 2023, 2020b), whereas the anterior temporal lobe (ATL) acts as an amodal hub of the conceptual system (Jefferies, 2013; Lambon Ralph et al., 2016; Patterson et al., 2007).

Crucially, the recruitment of both modality-specific and multimodal regions is task-dependent. Several studies indicate that modality-specific perceptual-motor regions are selectively engaged when the task requires the retrieval of perceptual-motor features of concepts (Binder and Desai, 2011; Hsu et al., 2011; Kemmerer, 2015; Willems and Casasanto, 2011). For example, we previously showed that auditory regions are selectively recruited for sound features during sound judgments, whereas somatomotor regions are selectively engaged for action features during action judgments (Kuhnke et al., 2021, 2020b; for similar results, see Borghesani et al., 2019; Hoenig et al., 2008; Popp et al., 2019a; van Dam et al., 2012). Remarkably, multimodal regions (e.g., left IPL and pMTG) also showed a task-dependent activation profile, responding to sound features during sound judgments and to action features during action judgments (Kuhnke et al., 2020b).

However, several issues are still open. First, it remains unclear which brain regions contain neural representations of perceptual-motor features of concepts, and to what extent these feature representations are also modulated by the task. Previous neuroimaging studies of conceptual feature retrieval have mainly investigated task-dependent changes in general recruitment of brain regions, that is, changes in mean activation magnitude via univariate analyses (Hsu et al., 2011; Kemmerer, 2015; Kuhnke et al., 2020b; van Dam et al., 2012). However, neural representations of mental contents are generally assumed to be encoded in “population codes”—patterns of activity distributed across multiple representational units (Connolly et al., 2012; Haxby et al., 2014; Ritchie et al., 2019). Crucially, some brain regions might only show task-dependent modulations of conceptual feature representations—as reflected in *relative* differences between fine-grained activity patterns—without a change in *absolute* engagement (Kriegeskorte and Bandettini, 2007; Raizada and Kriegeskorte, 2010). Whereas univariate analyses are insensitive to such fine-grained activity pattern differences, population codes can be studied using multivariate pattern analyses (MVPA) of functional neuroimaging data (Haxby, 2012; Mur et al., 2009). Therefore, MVPA might reveal task-dependent conceptual feature representations in brain regions that have remained undetected by univariate analyses. While several previous MVPA studies have shown task- or context-dependent modulations of activity patterns during conceptual processing (Aglinskas and Fairhall, 2023; Fu et al., 2023; Gao et al., 2022a, 2021; Liuzzi et al., 2021; Xu et al., 2018), potential task-dependent modulations of individual perceptual-motor feature representations have not been investigated yet.

Second, it is unknown whether conceptual representations in multimodal regions are indeed cross-modal, that is, similar for different modalities. As multimodal regions are typically identified via conjunctions of brain activation maps (Fernandino et al., 2016a; Kuhnke et al., 2020b), it is possible that multimodal overlap reflects spatially overlapping but distinct fine-grained activity patterns for different modalities (Downing et al., 2007; Haxby et al., 2001).

Third, it remains unclear how the conceptual system is related to the large-scale functional networks of the human brain, as identified using resting-state functional connectivity MRI (Buckner et al., 2009; Yeo et al., 2011). Several authors have noted the topographical similarity of the conceptual system, especially cross-modal areas, to the default mode network (DMN) (Binder et al., 2009, 1999; Fernandino et al., 2016a). The DMN is a set of brain regions that show deactivation during attention-demanding tasks (as compared to rest), and strong functional coupling during the resting state (Buckner et al., 2008; Raichle et al., 2001). The DMN is engaged in spontaneous thought, self-referential and autobiographical processes, as well as mentalizing (Andrews-Hanna, 2012; Smallwood et al., 2021). These forms of introspective information may contribute to conceptual knowledge (Kiefer et al., 2022; Ulrich et al., 2022). In addition, conceptual processing is frequently assumed to involve domain-general executive control or “multiple demand” systems, such as the frontoparietal control network (FPN) and/or the dorsal attention network (DAN) (Hodgson et al., 2021; Noonan et al., 2013; Wang et al., 2021). Specifically, FPN and DAN may support the controlled retrieval and/or selection of task-relevant conceptual representations (Noonan et al., 2013; Thompson-Schill et al., 1999; Wagner et al., 2001).

Here, we asked (1) which brain regions show task-dependent modulations of conceptual feature representations, (2) whether conceptual representations in putative multimodal regions are indeed cross-modal, and (3) how the brain regions engaged in conceptual feature retrieval relate to the large-scale functional brain networks. To this end, we conducted MVPA decoding analyses on our previous fMRI data (Kuhnke et al., 2021, 2020b). 40 participants performed three different tasks—lexical decision, sound judgment, and action judgment—on written words with a high or low association to sounds and actions (e.g., “telephone” is a high sound–high action word). Sound and action feature associations were completely crossed, enabling the independent investigation of sound and action feature representations.

First, in “searchlight” decoding analyses, we localized brain regions enabling above-chance decoding of sound and action features of concepts. In each task, we moved a spherical region-of-interest (or “searchlight”) through the entire brain and trained a machine-learning classifier to decode high vs. low action or sound words based on the activity patterns within each searchlight. We compared the results for searchlight MVPA to classical univariate analysis to identify additional information represented in fine-grained activity patterns. Next, to test for cross-modal representations of task-relevant conceptual features, we trained a classifier on sound features (high vs. low sound words) during sound judgments, and tested the classifier on action features (high vs. low action words) during action judgments, and vice versa. Finally, we investigated the involvement of the large-scale functional brain networks, as characterized in the resting-state network parcellation by Yeo et al. (2011). To this end, we assessed the spatial overlap between the MVPA searchlight maps and each functional network, and performed MVPA decoding analyses using the activity patterns within each network separately.

We hypothesized that conceptual representations are modulated by the task: In both modality-specific and multimodal brain regions, activity patterns for sound and action features should be most distinctive when they are task-relevant. Secondly, multimodal regions should contain cross-modal conceptual representations, enabling cross-decoding between task-relevant sound and action features. Finally, we expected that conceptual feature retrieval involves the DMN, and possibly domain-general control (FPN) and attention (DAN) networks.

## 2. Methods

### 2.1. Subjects

Data from 40 healthy native German speakers (22 female; mean age: 26.6 years; SD: 4.1; range: 19-33) were analyzed. 42 participants were initially recruited, but two were excluded due to strong head movement or aborting the experiment. All participants were right-handed (mean laterality quotient: 93.7; SD: 9.44; Oldfield, 1971) and had no history of neurological disorders or head injury, or exhibited contraindications to fMRI. They were recruited via the subject database of the Max Planck Institute for Human Cognitive and Brain Sciences, Leipzig, Germany. Written informed consent was obtained from each subject prior to the experiment. The study was performed according to the guidelines of the Declaration of Helsinki and approved by the local ethics committee of the University of Leipzig.

### 2.2. Experimental procedures

The experimental procedure is reported in detail in Kuhnke et al. (2020b), and summarized here. We used a 3 x 2 x 2 within-subject design with the factors TASK (lexical decision, sound judgment, action judgment), SOUND (high, low association), and ACTION (high, low association). In two event-related fMRI sessions, participants performed three different tasks—lexical decision, sound judgment, and action judgment—on 192 written words with a high or low association to sounds and actions (Figure 1). In the first session, participants performed a lexical decision task, in which they decided whether the presented stimulus was a real word or pseudoword. In the other session, participants performed the sound and action judgment tasks. In the sound judgment task, participants judged whether the object denoted by the word was strongly associated with sounds. In the action judgment task, participants judged whether the object was strongly associated with actions. Whereas the lexical decision task acted as an implicit control task that did not require sound or action knowledge, the sound and action judgment tasks explicitly required sound and action knowledge, respectively. Instructions were given at the beginning of each scanning session. Participants practiced all tasks outside the scanner before the session with 16 trials that were excluded from the main experiment.

**Figure 1.**
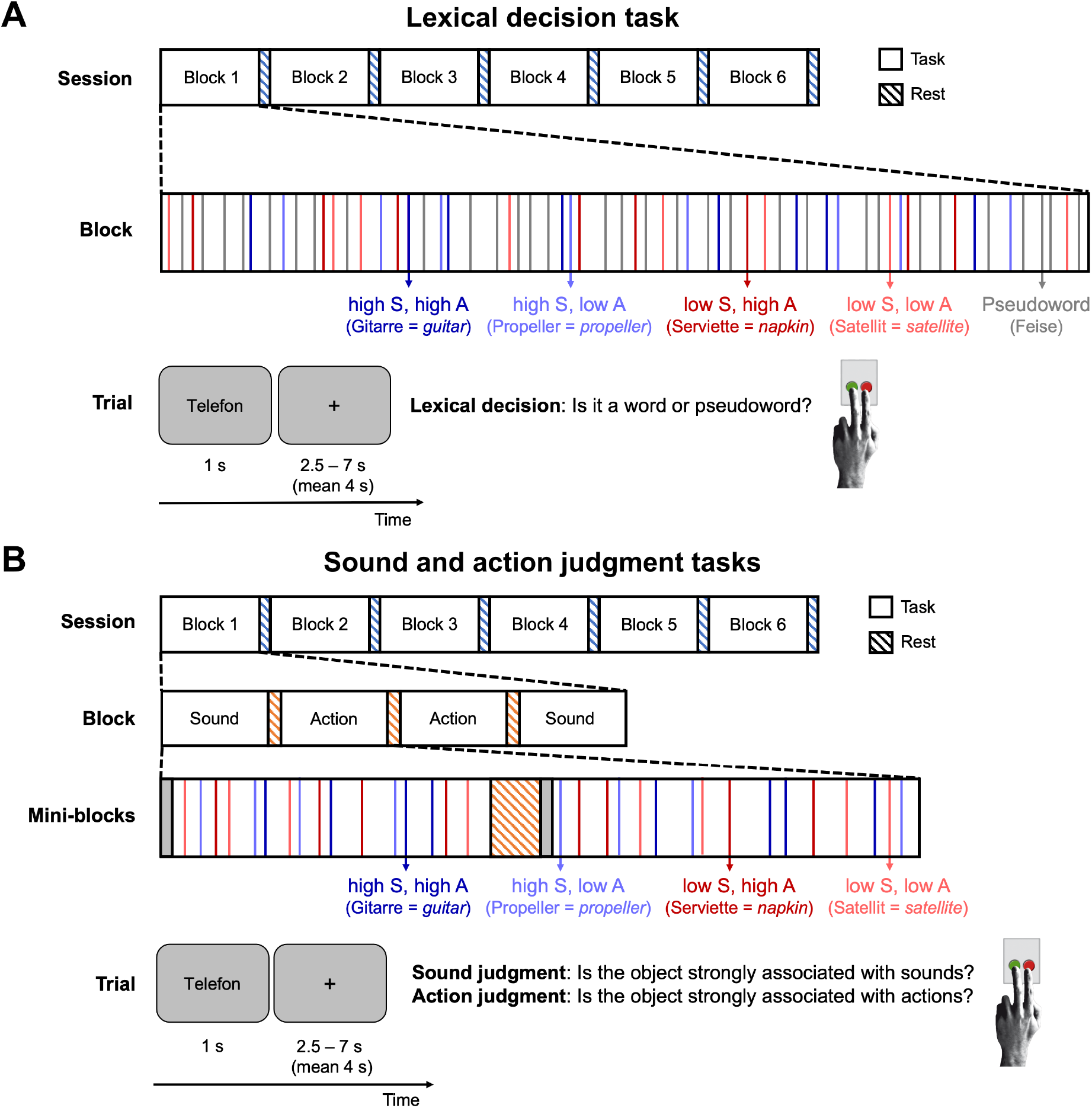
Experimental procedure. In two fMRI sessions, participants performed a lexical decision task (A), and sound and action judgment tasks (B). Trials for the four word types high sound–high action (dark blue), high sound–low action (light blue), low sound–high action (dark red), and low sound–low action (light red) were presented in random order within 6 blocks (64 trials each). Blocks were separated by 20-s rest periods (blue-striped bars). Sound and action judgment tasks were performed in mini-blocks of 16 trials, separated by 12-s rest periods (orange-striped bars). Each mini-block started with a cue indicating the task. On each trial, a word was shown for 1 s, followed by an inter-trial interval (fixation cross) of 2.5-7 s. Participants responded via button press. Adapted from Kuhnke et al. (2021, 2020b).

High and low sound words selectively differed in their association to sounds, while high and low action words selectively differed in their association to actions, as determined by the ratings of a different group of 163 volunteers (cf. Fernandino et al., 2016; Trumpp et al., 2014). Word types were matched on ratings of the respective other feature, visual conceptual associations and familiarity, number of letters and syllables, word frequency, bi- and trigram frequencies, and number of orthographic neighbors (Table S1). Stimuli for all conditions were selected from the same superordinate categories of animals, inanimate natural entities, and man-made objects (Goldberg et al., 2006; Kiefer et al., 2008). For the lexical decision task, a pseudoword was generated for each word matched in length, syllable structure and transition frequencies using the *Wuggy* software (Keuleers and Brysbaert, 2010; http://crr.ugent.be/Wuggy). For the full stimulus set, see the Supplementary Material of Kuhnke et al. (2020a).

### 2.3. fMRI acquisition and preprocessing

fMRI data were collected on a 3T Prisma scanner (Siemens, Erlangen, Germany) equipped with a 32-channel head coil. Functional blood oxygenation level dependent (BOLD) images were acquired using a multiband dual-echo EPI sequence (repetition time (TR): 2 s; echo times (TE): 12 & 33 ms; flip angle: 90°; field of view (FoV): 204 mm; voxel size: 2.5 x 2.5 x 2.5 mm; slice gap: 0.25 mm; bandwidth: 1966 Hz/Px; phase encoding direction: A/P; multiband factor 2). We used a dual-echo sequence (Halai et al., 2014; Poser et al., 2006) and tilted slices 10° up from the AC-PC line (Weiskopf et al., 2006) to minimize susceptibility artifacts and maximize BOLD sensitivity throughout the entire brain, including in regions suffering from signal loss in single-echo EPI such as the ATL (Devlin et al., 2000). B0 field maps were acquired for susceptibility distortion correction using a gradient-echo sequence (TR: 0.62 s; TE: 4 & 6.46 ms; flip angle: 60°; bandwidth: 412 Hz/Px; other parameters identical to functional sequence). Structural T1-weighted images were acquired for normalization using an MPRAGE sequence (176 slices in sagittal orientation; TR: 2.3 s; TE: 2.98 ms; FoV: 256 mm; voxel size: 1 × 1 × 1 mm; no slice gap; flip angle: 9°; phase encoding direction: A/P).

fMRI analysis was performed using *Statistical Parametric Mapping* (*SPM12*; Wellcome Trust Centre for Neuroimaging; http://www.fil.ion.ucl.ac.uk/spm/) implemented in *Matlab* (version 9.10). The two images with a short and long TE were combined using an average weighted by the temporal signal-to-noise ratio (tSNR) at each voxel, which yields optimal BOLD sensitivity (Poser et al., 2006). tSNR was calculated based on 30 volumes collected at the beginning of each scanning run, which were excluded from further analyses. Functional images were realigned, distortion corrected, slice-timing corrected, and normalized to MNI space (via normalization of the coregistered structural image).

### 2.4. Univariate analyses

Univariate analysis employed the classical two-level approach in SPM. At the first level, individual subject data smoothed with a 5 mm^3^ FWHM Gaussian kernel were modeled using the general linear model (GLM). The subject-level GLM included one regressor for each experimental condition (13 in total), modeling trials as stick functions convolved with the canonical HRF and its temporal derivative. Lexical decisions on pseudowords were modelled in the GLM, but excluded from further analyses. Only correct trials were analyzed, error trials were modeled in a separate regressor-of-no-interest. Nuisance regressors included the 6 motion parameters, individual regressors for time points with strong volume-to-volume movement (framewise displacement > 0.9; Siegel et al., 2014), and a duration-modulated parametric regressor accounting for response time differences between trials and conditions (Grinband et al., 2008). The data were subjected to an AR(1) auto-correlation model to account for temporal auto-correlations, and high-pass filtered (cutoff 128 s) to remove low-frequency noise.

Contrast images were computed at the first level for each participant. At the second (group) level, these contrast images were submitted to non-parametric permutation tests (5000 permutations; *SnPM toolbox*; https://warwick.ac.uk/fac/sci/statistics/staff/academic-research/nichols/software/snpm/). To identify brain regions sensitive to action or sound features of concepts in each task (lexical decisions, action judgments, sound judgments), we compared activation for high > low action words, and high > low sound words in each task. To localize “multimodal regions” engaged in both sound and action feature retrieval, we performed conjunction analyses between [sound judgment: high > low sound words] ∩ [action judgment: high > low action words] via minimum-statistic conjunctions (testing the conjunction null; Nichols et al., 2005). That is, a voxel was only considered significant in the conjunction if it was significant for both contrasts. All activation maps were thresholded at a voxel-wise p < 0.001 and a cluster-wise p < 0.05 FWE-corrected for multiple comparisons. Notably, to optimally match the univariate and MVPA decoding analyses, our current univariate analyses slightly differ from those in our previous publication (Kuhnke et al., 2020b) in smoothing (5 vs. 8 mm^3^) and thresholding (non-parametric cluster-wise FWE vs. voxel-wise FDR correction).

### 2.5. MVPA searchlight analyses

To allow for valid comparison to our univariate analyses, searchlight MVPA was performed with as similar parameters as possible. As for univariate analysis, individual subject data were modeled separately using the GLM. The subject-level GLM for MVPA was identical to the univariate GLM (i.e., same HRF model, same nuisance regressors, same auto-correlation model and high-pass filtering), with the following exceptions: First, MVPA was performed on unsmoothed subject-level data as is common for MVPA to retain fine-grained multi-voxel activity patterns (Haxby et al., 2014; Raizada and Lee, 2013). Second, the two sessions (lexical decision task vs. sound and action judgment tasks) were modelled in separate GLMs. Third, the subject-level GLMs for MVPA included one regressor for each trial to obtain trial-wise activity estimates. The GLM for lexical decisions included 384 regressors: 192 for words, and 192 for pseudowords. The GLM for sound and action judgments included 408 regressors: 192 for sound judgments, 192 for action judgments, and 24 for task cues.

Next, these subject-specific trial-wise estimates were used as input for MVPA decoding using *The Decoding Toolbox* (version 3.999E; Hebart et al., 2015) implemented in *Matlab* (version 9.10). For searchlight MVPA, we moved a spherical region-of-interest (or “searchlight”) of 5 mm radius (27 voxels in total) through the entire brain (Kriegeskorte et al., 2006). We first aimed to localize neural representations of action and sound features of concepts in each task. At each searchlight location, a machine-learning classifier (an L2-norm support vector machine; C=1) aimed to decode between high vs. low action words, as well as high vs. low sound words, within each task (yielding 6 different decoding analyses). We performed leave-one-block-out cross validation, training on the activity patterns for 5 blocks, and testing on the remaining 6^th^ block (Oosterhof et al., 2012; Varoquaux et al., 2017). Therefore, the classifier was trained and tested on independent sets of data (Hebart et al., 2015). Each block was used once for testing, which yielded 6 cross-validation iterations.

To identify cross-modal representations of task-relevant conceptual features, we performed “cross-decoding”, that is, training and testing the classifier on different experimental conditions (Skerry and Saxe, 2014; Wurm and Lingnau, 2015). Specifically, we trained the classifier on sound features (high vs. low sound words) in the sound judgment task, and tested the classifier on action features (high vs. low action words) in the action judgment task. We also performed training and testing in the reverse direction, and averaged the results for each subject before group analysis.

Subject-specific classification accuracy maps (minus chance accuracy of 50%) were smoothed with a 5 mm^3^ FWHM Gaussian kernel, matching the smoothing level for our univariate analyses. Finally, the smoothed subject-specific accuracy maps were entered into non-parametric permutation tests at the group level (5000 permutations; *SnPM toolbox*). The right somatomotor cortex was masked out to remove brain activity related to left-handed button presses (using a mask of right M1/S1/PMC/SMA from the human motor area template; Mayka et al., 2006). As for our univariate analyses, all MVPA searchlight maps were thresholded at a voxel-wise p < 0.001 and cluster-wise p < 0.05 FWE-corrected.

### 2.6. Comparison of univariate analysis and MVPA decoding in anatomical ROIs

In addition to our comparison at the whole-brain level, we also compared univariate analysis and MVPA decoding in anatomical regions-of-interest (ROIs). We selected 7 anatomical ROIs that are commonly implicated in general conceptual-semantic processing (Binder and Desai, 2011; Kuhnke et al., 2023; Lambon Ralph et al., 2016): the left anterior inferior frontal gyrus (aIFG), middle frontal gyrus (MFG), posterior middle temporal gyrus (pMTG), posterior inferior parietal lobe (pIPL), as well as the bilateral anterior temporal lobe (ATL), posterior cingulate cortex (PCC) / precuneus, and dorsomedial prefrontal cortex (dmPFC).

Following the procedure by Hoffman and Tamm (2020), anatomical ROIs were extracted from the Harvard-Oxford atlas distributed with FSL (Jenkinson et al., 2012; Smith et al., 2004), thresholded at 30% probability to belong to the respective region, and binarized: aIFG = pars triangularis of the inferior frontal gyrus; MFG = middle frontal gyrus; pMTG = posterior middle temporal gyrus; pIPL = angular gyrus / posterior supramarginal gyrus; ATL = temporal pole / anterior superior, middle, inferior temporal / anterior parahippocampal / anterior fusiform gyri; PCC / precuneus = posterior cingulate cortex / precuneus; dmPFC = superior frontal gyrus.

For univariate analysis, we extracted mean activation magnitudes (contrast values in arbitrary units) from each ROI. For MVPA decoding, we performed leave-one-block-out cross validation, as for our searchlight analyses: A machine-learning classifier was trained on the activation patterns in a given ROI for 5 out of the 6 blocks, and tested on the remaining block. At the group level, activation magnitudes and classification accuracies (minus chance level of 50%) for each ROI and condition were entered into one-sample t-tests. Moreover, we performed (two-tailed) paired t-tests for differences in activation magnitude / decoding accuracy between conditions for each ROI. P-values were corrected for multiple comparisons via Bonferroni correction for the number of ROIs.

### 2.7. Spatial relationship between conceptual processing and large-scale functional networks

To assess the spatial relationship between conceptual brain regions revealed by searchlight MVPA and large-scale functional brain networks, we tested the overlap of our MVPA searchlight maps with the resting-state networks by Yeo et al. (2011). This functional overlap analysis was performed with both the 7-network and 17-network parcellations by Yeo et al. (2011). Specifically, we computed the percentage of voxels in our MVPA searchlight maps for action and sound feature retrieval, as well as cross-modal areas that fell into each large-scale network. As a measure for above-chance contribution of a functional network, the percentage overlap was compared to a baseline of equal contribution of each network (7-network parcellation: 100 / 7 = 14.29%; 17-network parcellation: 100 / 17 = 5.88%) using *χ*^2^-tests, correcting for multiple comparisons using Bonferroni correction.

### 2.8. ROI-based MVPA decoding in large-scale functional networks

As a more direct test of the involvement of each large-scale functional network in conceptual feature representation, we also performed MVPA decoding in ROIs corresponding to each functional network by Yeo et al. (2011). Our main analyses focused on the 7-network parcellation; the 17-network parcellation was tested in supplementary analyses (see Supplementary Material). Subject-level ROI-based decoding employed the same methods as our searchlight analyses, with the exception that the activation pattern across all voxels of the network was used for classification (Mur et al., 2009). Thus, we performed two types of MVPA decoding analyses: First, we aimed to decode action features (high vs. low action words) and sound features (high vs. low sound words) within each task (using leave-one-block-out cross validation). Second, we aimed to identify cross-modal representations of task-relevant conceptual representations. To this end, we performed cross-decoding, training a classifier on action features (high vs. low action words) in the action judgment task, and testing on sound features (high vs. low sound words) in the sound judgment task, and vice versa. As for our searchlight analyses, the right somatomotor cortex was masked out of the network ROIs to remove button press related activity.

At the group level, classification accuracies for each ROI and condition were entered into one-sample t-tests (vs. chance level of 50%). Moreover, we performed (two-tailed) paired t-tests for differences in decoding accuracy between conditions for a given ROI, and between ROIs for a given condition. P-values were corrected for multiple comparisons via Bonferroni correction for the number of ROIs.

## 3. Results

### 3.1. Whole-brain analyses: Univariate vs. MVPA

For both univariate analysis and searchlight MVPA, brain activity for action and sound features was strongly task-dependent.

#### 3.1.1. Action feature retrieval

During lexical decisions, neither univariate analysis nor MVPA revealed any significant brain activity for action features (high vs. low action words). During sound judgments, both univariate analysis and MVPA yielded activity for action features selectively in the left inferior parietal lobe (IPL), and no other brain region (Figure S1 A-C).

During action judgments, however, both univariate analysis and MVPA yielded widespread activity for action features. Univariate analysis (Figure 2A; Table S2) revealed action-related activations in left anterior inferior frontal gyrus (aIFG), IPL / intraparietal sulcus (IPS), posterior middle and inferior temporal gyri (pMTG/ITG), posterior cingulate cortex (PCC), caudate, and cerebellum.

**Figure 2.**
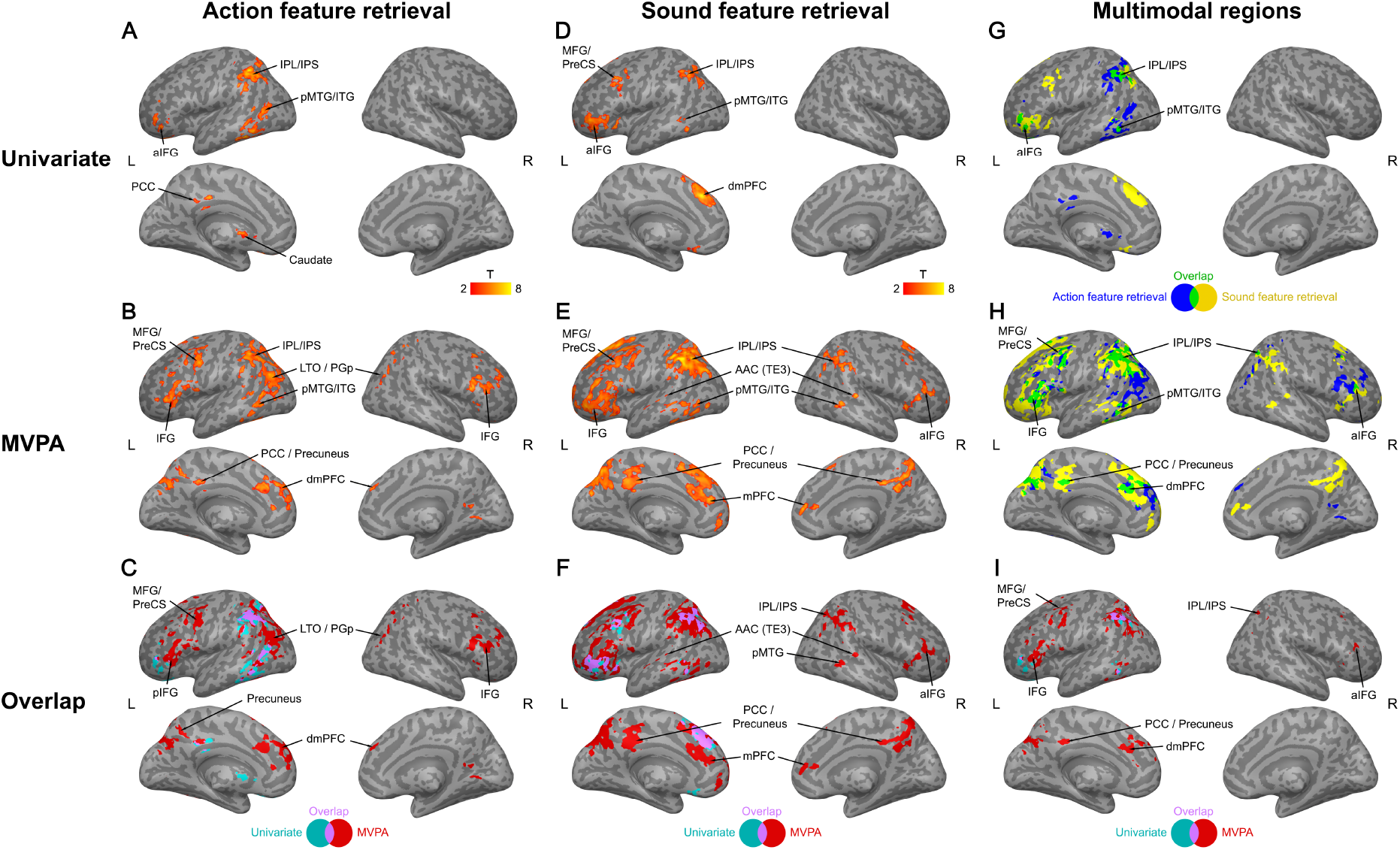
Comparison of results for whole-brain univariate analysis vs. searchlight MVPA on task-relevant conceptual feature retrieval. Both univariate and MVPA subject-specific maps were smoothed with a 5 mm^3^ FWHM Gaussian kernel. All group-level maps were thresholded at a voxel-wise p < 0.001 and a cluster-wise p < 0.05 FWE-corrected using non-parametric permutation tests.

Searchlight MVPA (Figure 2B; Table S3) found action-related activity in left IFG, IPL/IPS, pMTG/ITG, PCC / precuneus, the lateral temporo-occipital junction (LTO), middle frontal gyrus (MFG) / precentral sulcus (PreCS), dorsomedial prefrontal cortex (dmPFC), and cerebellum.

Comparison between univariate analysis and searchlight MVPA revealed overlap in left IPL/IPS, aIFG, pMTG/ITG, PCC, and cerebellum (Figure 2C purple; Table S4). However, MVPA activity patterns were broader in these regions, and only MVPA revealed recruitment of the right cerebral hemisphere, specifically in right IPL (area PGp), IFG, and LTO (Figure 2C red). Moreover, only MVPA found action-related activity in left posterior IFG, PMC, LTO, precuneus, and dmPFC.

#### 3.1.2. Sound feature retrieval

During lexical decisions, neither univariate analysis nor MVPA revealed any significant brain activity for sound features (high vs. low sound words). During action judgments, small clusters emerged in left IPL and bilateral PCC / precuneus, and no other area (Figure S1 D-F).

During sound judgments, both univariate analysis and MVPA revealed widespread activity for sound features. Univariate analysis (Figure 2D; Table S5) showed sound-related activations in left pMTG/ITG, IPL/IPS, aIFG, MFG/PreCS, dmPFC, and right cerebellum.

Searchlight MVPA (Figure 2E; Table S6) detected sound-related activity in bilateral IPL/IPS, pMTG/ITG, IFG, MFG/PreCS, mPFC, PCC / precuneus, cerebellum, as well as auditory association cortex (AAC; area TE3).

Comparison between univariate analysis and searchlight MVPA showed overlap in left IPL/IPS, aIFG, pMTG/ITG, and dmPFC (Figure 2F purple; Table S7). However, MVPA activity patterns were more extensive in these areas, and only MVPA revealed engagement of the right cerebral hemisphere, specifically in right IPL/IPS, pMTG, and aIFG (Figure 2F red). Moreover, only MVPA revealed sound-related activity in bilateral PCC / precuneus, mPFC, and AAC (area TE3).

#### 3.1.3. Multimodal regions

To identify multimodal regions engaged in both action and sound feature retrieval, we performed conjunction analyses between [action judgments: high vs. low action words] and [sound judgments: high vs. low sound words] for both univariate analyses and searchlight MVPA.

Univariate analysis (Figure 2G; Table S8) identified multimodal regions in left IPL/IPS, pMTG/ITG, aIFG, and right cerebellum. Searchlight MVPA (Figure 2H; Table S9) found multimodal regions in left pMTG/ITG, MFG/PreCS, dmPFC, PCC / precuneus, cerebellum, as well as in bilateral IPL/IPS and IFG.

Direct comparison between univariate analysis and MVPA revealed overlap in left IPL/IPS, pMTG/ITG, and aIFG (Figure 2I purple; Table S10). However, MVPA multimodal clusters were broader in all of these areas, and only MVPA yielded multimodal regions in the right cerebral hemisphere, specifically in right IPL/IPS, and aIFG (Figure 2I red). In addition, only MVPA revealed multimodal overlap in left MFG/PreCS, PCC / precuneus and dmPFC.

#### 3.1.4. Cross-modal conceptual representations

Next, we assessed whether “multimodal” overlap between sound and action feature retrieval was indeed based on cross-modal conceptual representations. As the machine-learning classifier was trained and tested on sound and action features separately, it is possible that successful decoding relied on spatially overlapping but distinct activity patterns. We reasoned that if these regions indeed hold cross-modal conceptual representations, it should be possible to train a classifier on sound features during sound judgments, and test it on action features during action judgments, and vice versa.

We found that cross-decoding of task-relevant conceptual features was possible (i.e., significant above chance level) in bilateral IPL/IPS, as well as in left precuneus and dmPFC (Figure 3; Table S11).

**Figure 3.**
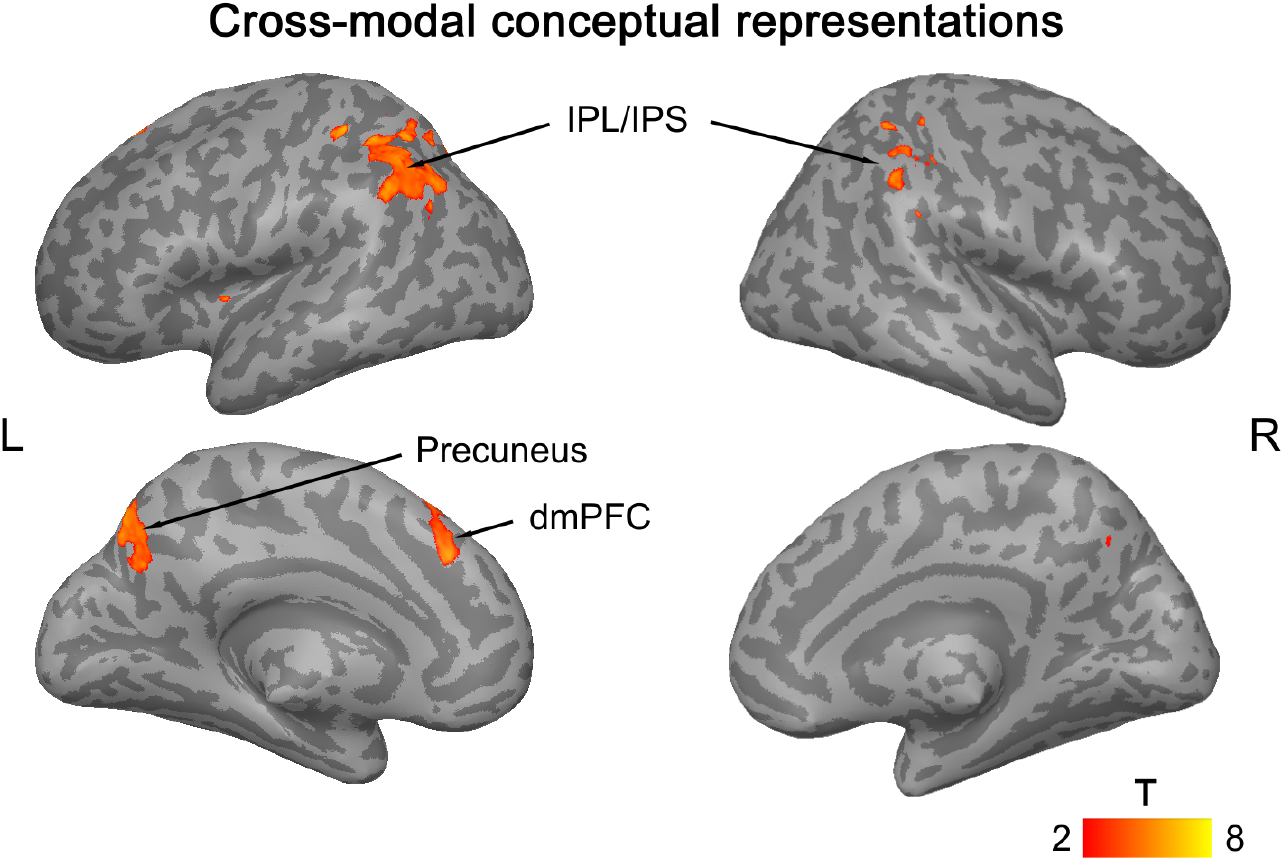
Brain regions showing significant cross-decoding of task-relevant conceptual features. The classifier was trained on activation patterns for task-relevant sound features (sound judgments: high vs. low sound words) and tested on task-relevant action features (action judgments: high vs. low action words), and vice versa. The searchlight map was thresholded at a voxel-wise p < 0.001 and a cluster-wise p < 0.05 FWE-corrected using non-parametric permutation tests.

### 3.2. ROI Analyses: Univariate vs. MVPA

In addition to our whole-brain analyses, we also compared MVPA decoding and univariate analysis in anatomical regions-of-interest (ROIs) commonly implicated in conceptual-semantic processing: left aIFG, MFG, pMTG, pIPL, as well as bilateral ATL, PCC / precuneus and dmPFC. The results further support the increased sensitivity of MVPA to task-dependent conceptual feature representations (Figure 4).

**Figure 4.**
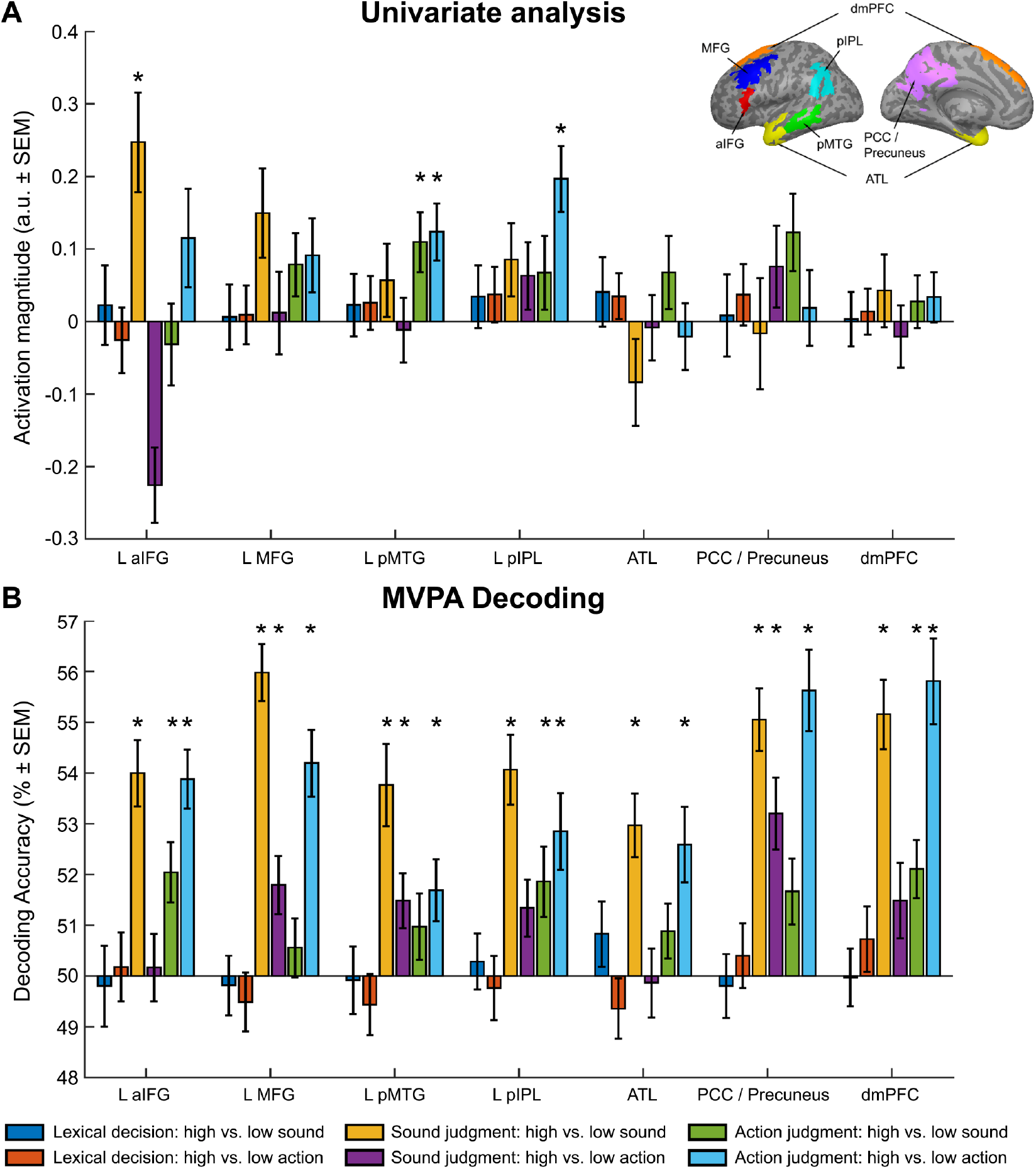
Comparison of univariate analysis and MVPA decoding in anatomical ROIs. (A) For univariate analysis, we extracted mean activation magnitudes (contrast values in arbitrary units; a.u.) from each ROI. (B) For MVPA decoding, a machine-learning classifier was trained on the activation patterns in a given ROI for 5 out of the 6 blocks, and tested on the remaining block (i.e., leave-one-block-out cross validation). *: p < 0.05 (Bonferroni-corrected for the number of ROIs).

Univariate analysis (Figure 4A; Tables S12-13) revealed significant activations for sound features (high vs. low action words) during sound judgments in left aIFG, and during action judgments in left pMTG. Action features (high vs. low action words) induced significant activity selectively during action judgments in left pMTG and pIPL. Left MFG, ATL, PCC / precuneus, and dmPFC did not show significant activations in this analysis.

In contrast, MVPA decoding (Figure 4B; Table S14) revealed significant activity for task-*relevant* conceptual features in all anatomical ROIs: for sound features (high vs. low sound words) during sound judgments, and for action features (high vs. low action words) during action judgments. All regions except ATL also showed some activity for task-*irrelevant* features: for sound features during action judgments in left aIFG, pIPL, and dmPFC; and for action features during sound judgments in left MFG, pMTG, and PCC / precuneus. However, decoding accuracies were generally higher for task-relevant than –irrelevant features (Table S15).

### 3.3. Relationship between conceptual processing and large-scale functional networks

To investigate the relationship between brain regions engaged during conceptual feature retrieval and the large-scale functional networks of the human brain, we analyzed the overlap between our MVPA searchlight maps and the resting-state networks by Yeo et al. (2011).

#### 3.3.1. Action feature retrieval

In the 7-network parcellation, action feature retrieval (action judgments: high vs. low action words) mainly involved parts of the default (27.6% voxels; χ^2^ = 335.13, p < 0.001), frontoparietal control (24.9%; χ^2^ = 223.20, p < 0.001), and dorsal attention (18.4%; χ^2^ = 38.93, p < 0.001) networks (Figure 5A and C).

In the 17-network parcellation, action feature retrieval overlapped with the dorsal attention A (13.5%; χ^2^ = 207.01, p < 0.001), control B (17.5%; χ^2^ = 407.36, p < 0.001) and control C (9.8%; χ^2^ = 67.36, p < 0.001), as well as the default C (10.6%; χ^2^ = 91.45, p < 0.001) and temporo-parietal (12.4%; χ^2^ = 160.16, p < 0.001) networks (Figure 5B).

**Figure 5.**
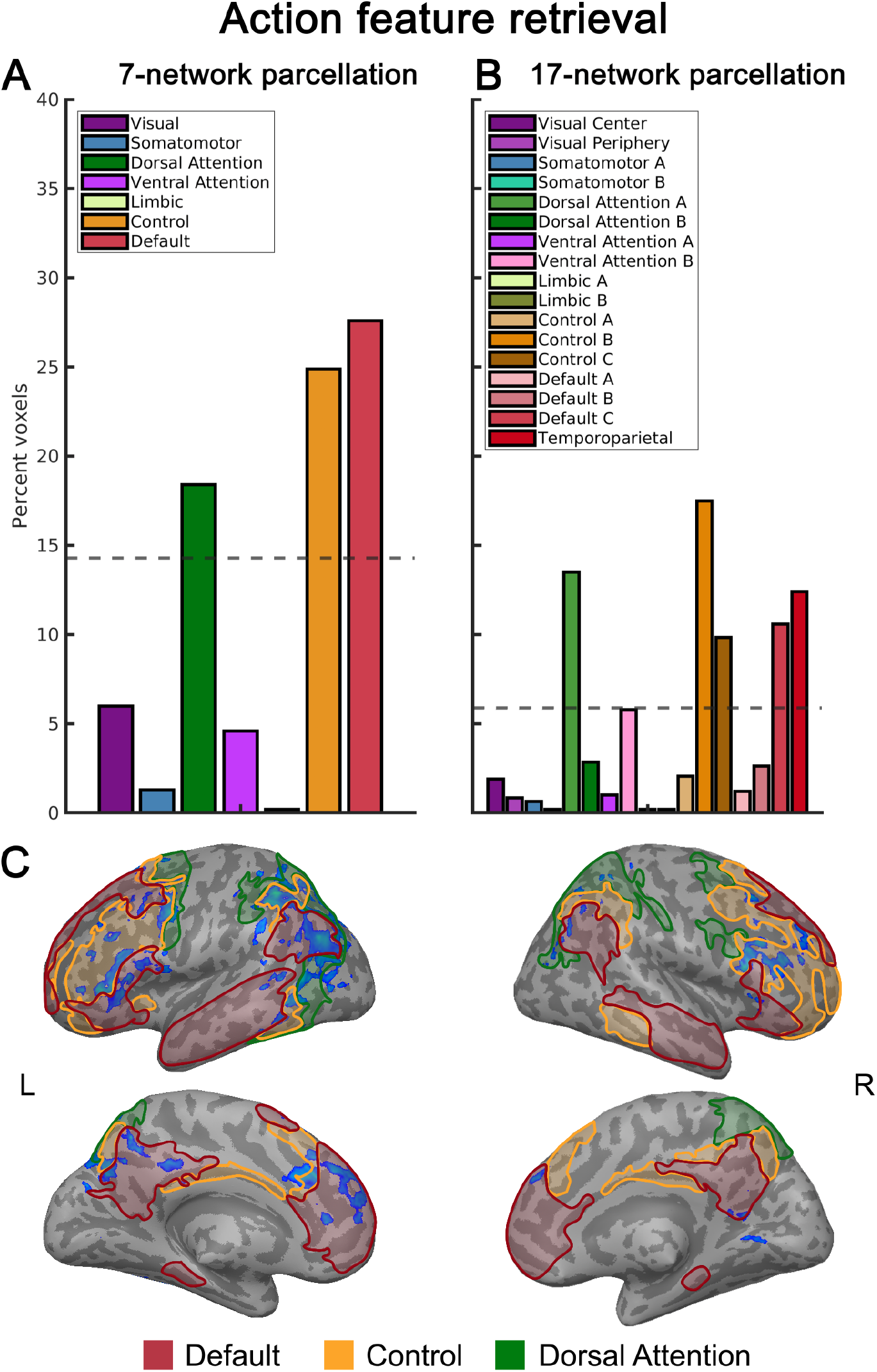
Overlap between the MVPA searchlight map for action feature retrieval and the resting-state networks by Yeo et al. (2011). We investigated both the 7-network (A) and 17-network (B) parcellations. Dashed lines represent the baseline level of equal overlap with each network. (C) Illustration of the three core networks from the 7-network parcellation that overlap with the MVPA searchlight map for action feature retrieval (blue; action judgments: high vs. low action words).

#### 3.3.2. Sound feature retrieval

In the 7-network parcellation, sound feature retrieval (sound judgments: high vs. low sound words) mainly involved the default (34.9% voxels; χ^2^ = 335.13, p < 0.001) and frontoparietal control (26.7%; χ^2^ = 223.20, p < 0.001) networks (Figure 6A and C). The dorsal attention network also showed some overlap (13.1%), but below the baseline level of equal overlap with each network (14.29%).

**Figure 6.**
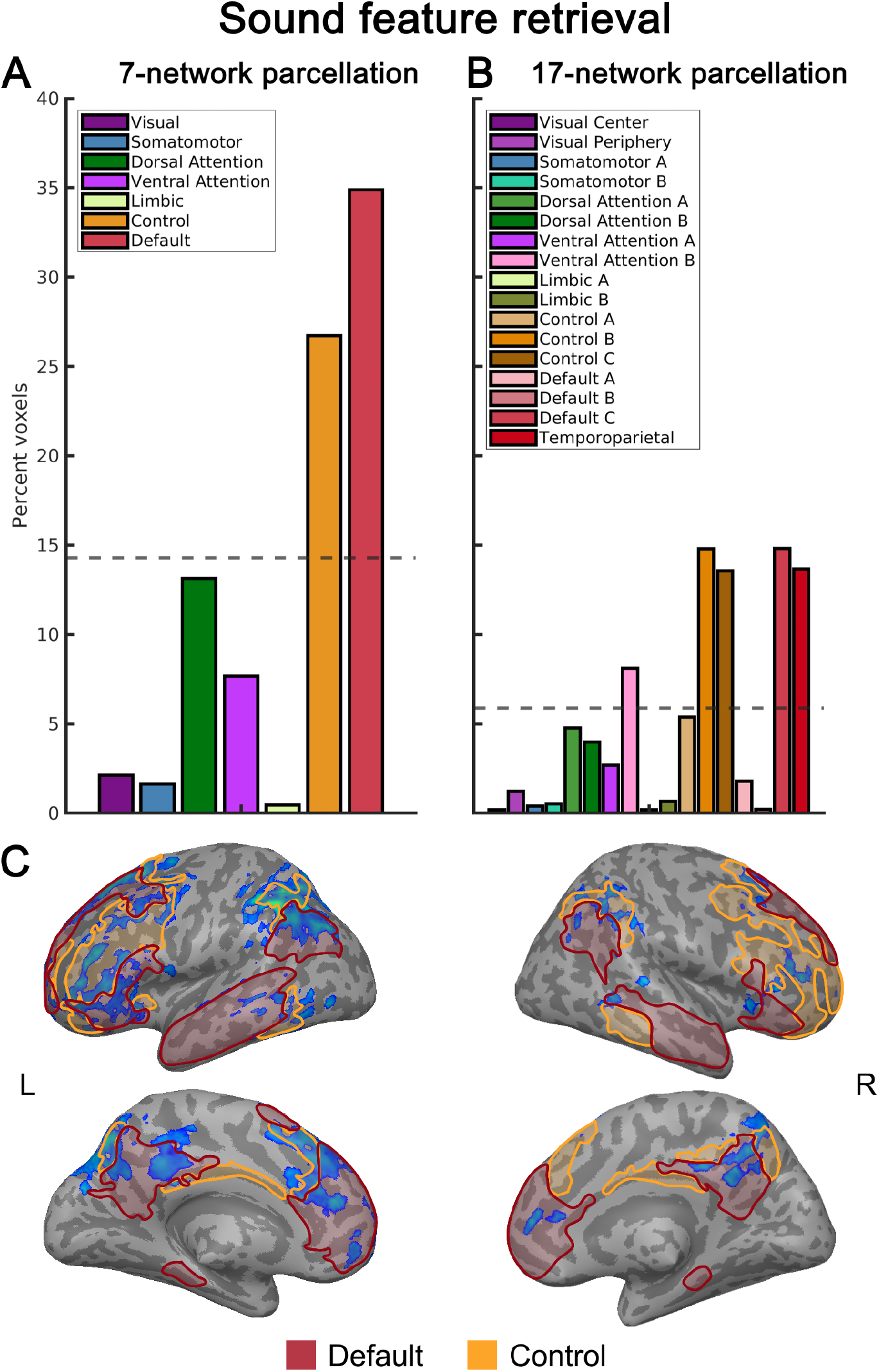
Overlap between the MVPA searchlight map for sound feature retrieval and the resting-state networks by Yeo et al. (2011). We investigated both the 7-network (A) and 17-network (B) parcellations. Dashed lines represent the baseline level of equal overlap with each network. (C) Illustration of the two core networks from the 7-network parcellation that overlap with the MVPA searchlight map for sound feature retrieval (blue; sound judgments: high vs. low sound words).

In the 17-network parcellation, sound feature retrieval overlapped with the control B (14.8%; χ^2^ = 473.58, p < 0.001) and control C (13.6%; χ^2^ = 371.55, p < 0.001), default C (14.8%; χ^2^ = 474.36, p < 0.001) and temporo-parietal (13.7%; χ^2^ = 379.51, p < 0.001), as well as the saliency ventral attention B (8.1%; χ^2^ = 42.41, p < 0.001) networks (Figure 6B).

#### 3.3.3. Cross-modal conceptual representations

Finally, we compared the searchlight MVPA map for cross-decoding of task-relevant conceptual features to the resting-state networks. In the 7-network parcellation, cross-modal decoding mainly overlapped with the default (31.67%; χ^2^ = 125.94, p < 0.001), frontoparietal control (35.38%; χ^2^ = 175.99, p < 0.001) and dorsal attention (19.72%; χ^2^ = 15.30, p < 0.001) networks (Figure 7A and C).

**Figure 7.**
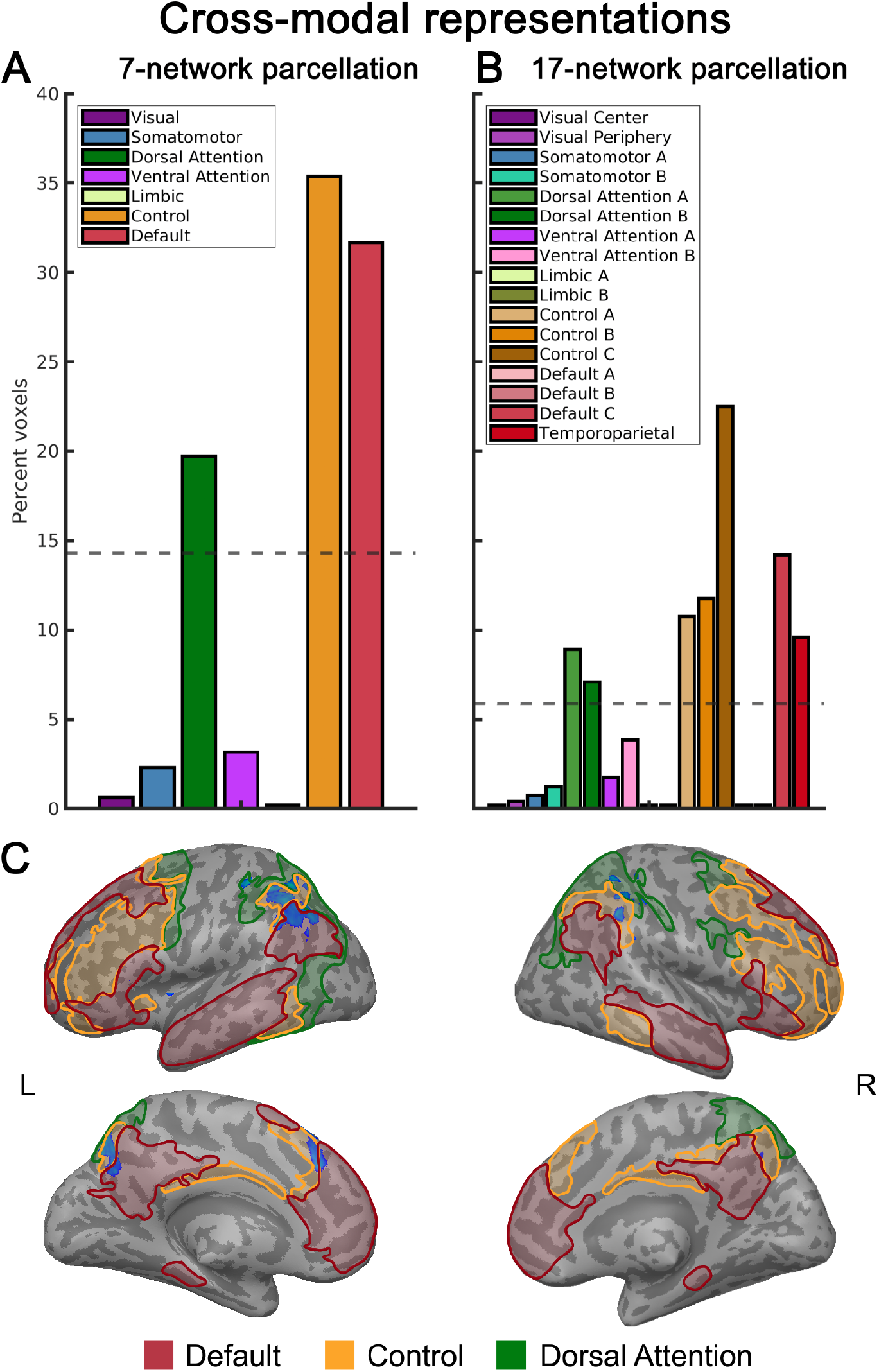
Overlap between the MVPA searchlight map for cross-modal conceptual representations and the resting-state networks by Yeo et al. (2011). We investigated both the 7-network (A) and 17-network (B) parcellations. Dashed lines represent the baseline level of equal overlap with each network. (C) Illustration of the three core networks from the 7-network parcellation that overlap with the MVPA searchlight map for cross-decoding (blue).

In the 17-network parcellation, cross-decoding overlapped with the control A (10.74%; X^2^ = 22.98, p < 0.001), control B (11.75%; X^2^ = 31.80, p < 0.001) and control C (22.38%; X^2^ = 167.89, p < 0.001), default C (14.18%; X^2^ = 56.62, p < 0.001) and temporo-parietal (9.59%; X^2^ = 14.32, p = 0.003), and dorsal attention A (8.91%; X^2^ = 9.98, p = 0.03) networks (Figure 7B). Involvement of the dorsal attention B network was not significantly above baseline level of equal overlap with each network (5.88%) (7.09%; X^2^ = 1.80, p > 0.05).

### 3.4. MVPA decoding in large-scale functional networks

As a more direct test of the involvement of large-scale functional networks in conceptual processing, we performed MVPA decoding based on the activation patterns in each network of the 7-network parcellation by Yeo et al. (2011).

Mirroring our searchlight analyses, we found that neural representations for sound and action features were strongly task-dependent (Figure 8; see Table S16 for statistics). During lexical decisions, no network displayed above-chance decoding of sound features (high vs. low sound words) or action features (high vs. low action words).

**Figure 8.**
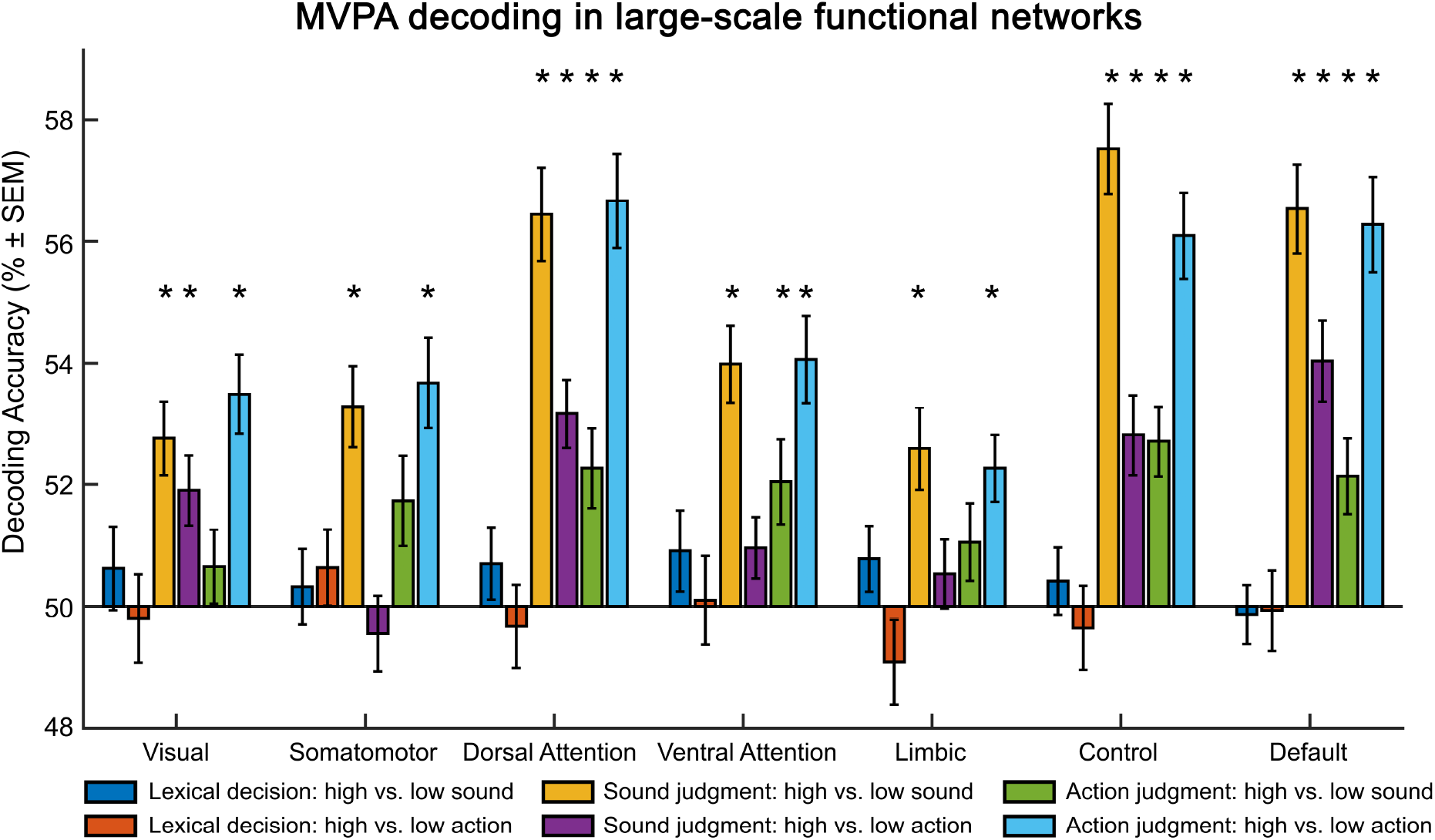
Results of ROI-based MVPA decoding analyses in the 7 resting-state networks by Yeo et al. (2011). A machine-learning classifier was trained on the activation patterns in a given network for 5 out of the 6 blocks, and tested on the remaining block (i.e., leave-one-block-out cross validation). *: p < 0.05 (Bonferroni-corrected for the number of networks).

In contrast, all networks showed significant decoding of task-*relevant* conceptual features: sound features during sound judgments (Figure 8 yellow), and action features during action judgments (Figure 8 cyan). However, decoding accuracies for task-relevant features were higher in the default, frontoparietal control, and dorsal attention networks than in the other networks (visual, somatomotor, saliency ventral attention, limbic) (Table S17).

Moreover, selectively the default, frontoparietal control, and dorsal attention networks enabled above-chance decoding of task-*irrelevant* conceptual features in both judgment tasks: action features during sound judgments (Figure 8 purple), and sound features during action judgments (Figure 8 green). Nonetheless, in all three networks, decoding accuracies were higher for task-relevant than -irrelevant features (Table S18).

The 17-network parcellation yielded similar results at a higher granularity (Figure S2).

#### 3.4.1. Cross-modal representations in large-scale functional networks

Finally, we also performed cross-decoding of task-relevant conceptual features in each network, training the classifier on sound features (high vs. low sound words) during sound judgments, and testing on action features (high vs. low action words) during action judgments, and vice versa. We found that cross-decoding was significant above chance in all networks except the limbic network (Figure 9; Table S19).

**Figure 9.**
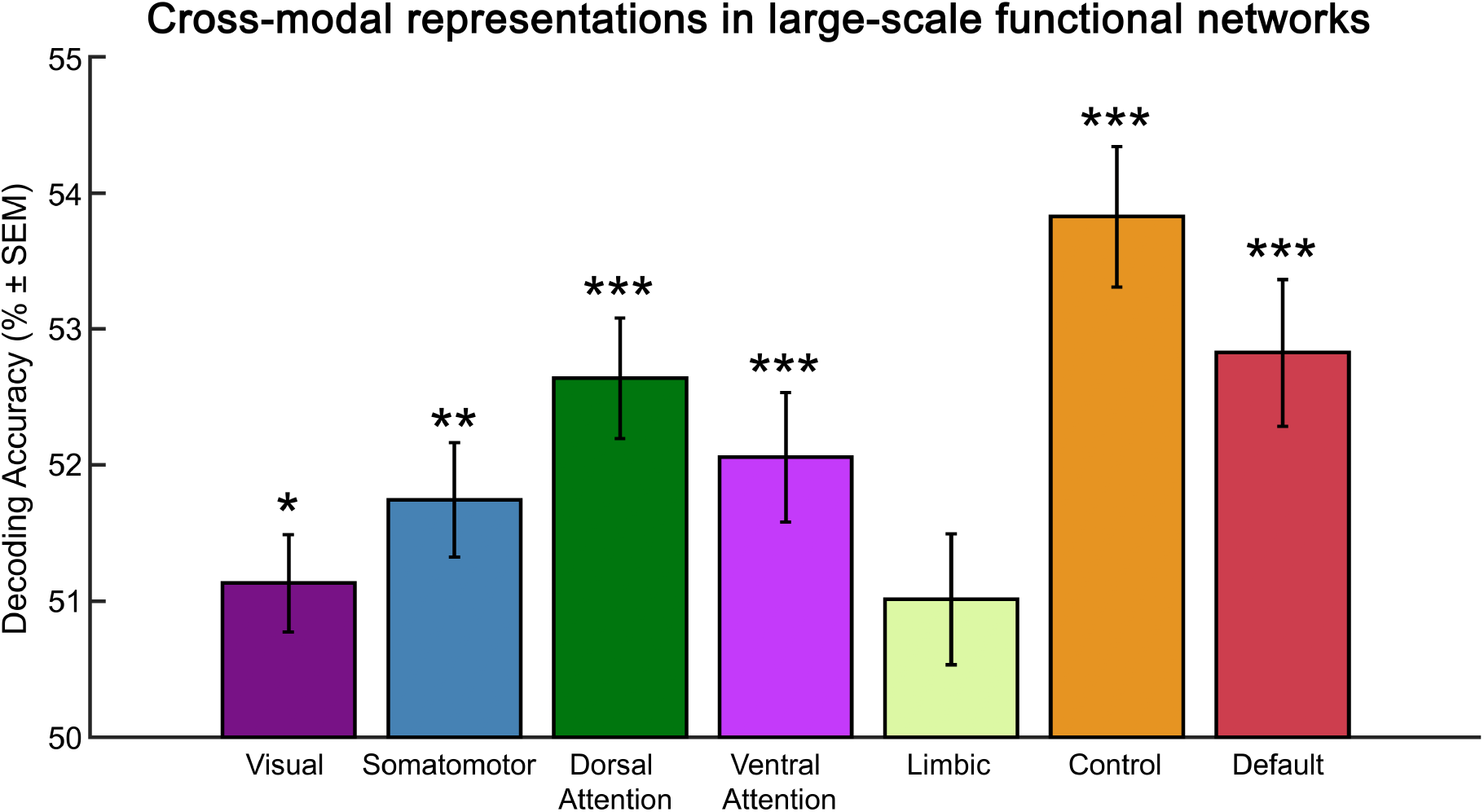
Cross-decoding of task-relevant conceptual features in the 7 resting-state networks by Yeo et al. (2011). The classifier was trained on activation patterns for task-relevant sound features (sound judgments: high vs. low sound words) and tested on task-relevant action features (action judgments: high vs. low action words), and vice versa. ***: p < 0.001; **: p < 0.01; *: p < 0.05 (Bonferroni-corrected for the number of networks).

However, decoding accuracies were higher in the frontoparietal control network than in the visual, somatomotor, salience ventral attention, limbic, and default networks (Table S20). Accuracies were higher in the default than visual and limbic networks; and decoding was more accurate in the dorsal attention than visual network.

Again, the 17-network parcellation showed similar results at a higher granularity (Figure S3).

## 4. Discussion

This study investigated neural representations of conceptual features in the human brain. Specifically, we asked (1) whether conceptual feature representations in modality-specific perceptual-motor and multimodal brain regions are modulated by the task, (2) whether conceptual representations in putative multimodal regions are indeed cross-modal, and (3) how the conceptual system relates to the large-scale functional brain networks.

We found that neural representations of conceptual features are strongly modulated by the task. Searchlight MVPA revealed task-dependent modulations of activity patterns for sound and action features of concepts: Both in modality-specific perceptual-motor and multimodal brain regions, activity patterns were most distinctive for sound features during sound judgments, and for action features during action judgments.

Several multimodal regions indeed showed evidence for cross-modal conceptual representations. Specifically, the bilateral inferior parietal lobe (IPL) / intraparietal sulcus (IPS), left precuneus and left dorsomedial prefrontal cortex (dmPFC) enabled cross-decoding of task-relevant conceptual features—from task-relevant sound to action features, and vice versa.

Finally, conceptual feature retrieval mainly involved the default mode network (DMN), frontoparietal control network (FPN), and dorsal attention network (DAN). MVPA searchlight maps for action and sound feature retrieval, as well as cross-modal areas showed extensive spatial overlap with these three networks. Direct MVPA decoding analyses within each network revealed that the DMN, FPN and DAN display the highest decoding accuracies for task-relevant conceptual features, constitute the only networks that could decode task-irrelevant features, and enable cross-decoding between task-relevant sound and action features.

These results suggest that conceptual representations in large-scale functional brain networks are task-dependent and cross-modal. Our findings support theories that assume conceptual processing to rely on a flexible, multi-level neural architecture.

### 4.1. Task dependency of conceptual representations

Our results indicate that conceptual feature representations encoded in fine-grained, multi-voxel activity patterns are strongly modulated by the concurrent task. Searchlight MVPA revealed by far the most extensive brain activity for sound and action features when they were task-relevant (i.e., for sound features during sound judgments, and for action features during action judgments).

These findings extend previous results from univariate neuroimaging analyses showing that the *general involvement* of brain regions in conceptual processing is task-dependent, with the strongest activation for task-relevant conceptual features (Borghesani et al., 2019; Hoenig et al., 2008; Hsu et al., 2011; Kemmerer, 2015; Kiefer and Pulvermüller, 2012; van Dam et al., 2012). For example, we previously found that both modality-specific perceptual-motor and multimodal brain regions are selectively engaged for sound features during sound judgments, and for action features during action judgments (Kuhnke et al., 2021, 2020b).

Compared to univariate analysis, MVPA revealed more extensive brain activity for task-relevant features in both modality-specific and multimodal regions. Only for MVPA, sound feature retrieval recruited the bilateral auditory association cortex (AAC) (Fernandino et al., 2016a; Kiefer et al., 2008; Trumpp et al., 2013a), while action feature retrieval engaged the bilateral lateral temporo-occipital junction (LTO) (Lewis, 2006; Oosterhof et al., 2010). These regions were selectively engaged for one feature, indicating that they are modality-specific (Barsalou, 2016; Kiefer and Pulvermüller, 2012). Regarding multimodal regions, MVPA decoding revealed more spatially extended multimodal effects in left anterior inferior frontal gyrus (aIFG), posterior middle and inferior temporal gyri (pMTG/ITG) and IPL/IPS than univariate analysis. Moreover, MVPA selectively revealed additional multimodal regions in left middle frontal gyrus (MFG) / precentral sulcus (PreCS), posterior cingulate cortex (PCC) / precuneus, dmPFC, as well as in right IPL/IPS and aIFG. These results converge with two recent MVPA studies demonstrating multimodal conceptual effects in bilateral IFG, MFG/PreCS, PCC / precuneus and dmPFC (Fernandino et al., 2022; Tong et al., 2022). Notably, whereas the precentral gyrus (pre- and primary motor cortex) seems to be specialized for action knowledge (Hauk et al., 2004; Kuhnke et al., 2023; Pulvermüller, 2013), our current results suggest that the adjacent precentral sulcus (PreCS) is multimodal.

Our whole-brain results are supported by a comparison of univariate analysis and MVPA decoding in anatomical ROIs: Only MVPA, but not univariate analysis, revealed multimodal feature representations in left aIFG, MFG, pMTG, pIPL, as well as bilateral ATL, PCC / precuneus, and dmPFC. In all of these regions, decoding accuracies were higher for task-relevant than –irrelevant features, indicating an enhancement of task-relevant conceptual feature representations.

Finally, only MVPA revealed activity related to conceptual feature representations in the right cerebral hemisphere. These findings suggest that the right hemisphere is also involved in conceptual processing, but plays a weaker role than the left hemisphere, at least under “normal” conditions in young and healthy human adults. In support of this view, Jung-Beeman (2005) summarized evidence that both the left and right hemispheres are engaged in conceptual-semantic cognition, but the right hemisphere seems to perform coarser computations than the left. This view is also corroborated by a recent large-scale fMRI study (n = 172) which revealed conceptual effects in both the left and right IPL, but stronger in the left (Kuhnke, Chapman et al., 2022).

Importantly, the task dependency of conceptual representations seems to be *graded*, rather than binary: Whereas no brain region showed significant activity for sound or action features during lexical decisions, we found some activity for sound features during action judgments, and for action features during sound judgments. These results suggest that in contrast to implicit tasks (e.g., lexical decision) which did not elicit feature-specific activity, explicit conceptual tasks (e.g., sound or action judgment) can induce task-*irrelevant* feature activation. In the present study, this effect might be explained by “action–sound coupling”, the phenomenon that many human actions are associated with typical sounds (e.g. hammering, guitar playing) (Lemaitre et al., 2018). Therefore, retrieval of action features of concepts may lead to the concomitant activation of associated sound features, and vice versa (Lemaitre et al., 2018; Lewis et al., 2006, 2005). However, this feature-related activity was restricted to high-level, cross-modal regions (e.g., left IPL, PCC / precuneus). Modality-specific perceptual-motor regions were selectively engaged when the respective feature was task-relevant (e.g., AAC during sound judgments, LTO during action judgments). These results are in line with the view that the recruitment of modality-specific perceptual-motor areas is particularly task-dependent (Binder and Desai, 2011; Kemmerer, 2015; Kuhnke et al., 2023; Willems and Casasanto, 2011). This view is now supported by several functional neuroimaging studies (Hoenig et al., 2008; Hsu et al., 2011; Kuhnke et al., 2021, 2020b; van Dam et al., 2012).

Notably, some studies found modality-specific activity even during shallow tasks, that is, implicit (Kiefer et al., 2012, 2008; Pulvermüller et al., 2005; Sim et al., 2015) or passive tasks (Hauk et al., 2004; Hauk and Pulvermüller, 2004), or when the stimulus was unattended (Pulvermüller and Shtyrov, 2006; Shtyrov et al., 2004) or not consciously perceived (Trumpp et al., 2014, 2013b). However, such effects have largely been observed when the pertinent feature was central to the concept. For instance, action verbs (e.g. “lick”, “kick”, or “pick”) engaged the motor cortex during shallow tasks (Hauk et al., 2004; Hauk and Pulvermüller, 2004; Tettamanti et al., 2005). As action knowledge is crucial to the meaning of action verbs, activation of motor regions might be required even for shallow comprehension of action verbs. These findings are thus consistent with the view that perceptual-motor features are selectively activated when relevant in the current context (cf. Kuhnke et al., 2020b).

Moreover, some studies showed that the semantic category structure between word meanings can be encoded in the neural similarity structure within modality-specific and multimodal regions, even in the absence of an explicit semantic task (e.g. during silent reading; Carota et al., 2021a, 2021b, 2017). For instance, semantic similarity between action-related verbs and nouns is encoded in left IFG, pMTG and premotor cortex, whereas left pITG represents semantic similarity between object-related nouns (Carota et al., 2017). These findings suggest that category structure between concepts can be encoded implicitly, whereas our results suggest that representations of individual perceptual-motor features of concepts are task-dependent, at least if they are not highly central for a given concept. Moreover, even if category representations can be detected in implicit tasks, they could be enhanced when task-relevant (Liuzzi et al., 2021; Xu et al., 2018). Relatedly, the sound features of different categories of objects may be represented differently in the human brain. For instance, Engel et al. (2009) demonstrated that human-produced, animal, mechanical, and environmental sounds preferentially engage different frontal and temporo-parietal structures, with an anterior-to-posterior division for living vs. non-living objects. While we controlled for such category effects in our stimulus design, future studies should directly investigate potential differences in sound feature representation for different categories of concepts (Trumpp et al., 2013a). Following action–sound coupling, sound features of action-related objects (e.g. the sound feature of “hammer”) might be tightly linked to both the auditory and somatomotor systems of the human brain (Lemaitre et al., 2018; Lewis et al., 2006, 2005).

Our study extends previous MVPA studies that demonstrated task- or context-dependent modulations of activity patterns during conceptual processing (Aglinskas and Fairhall, 2023; Fu et al., 2023; Gao et al., 2022a, 2021; Liuzzi et al., 2021; Xu et al., 2018). For instance, Gao et al. (2021) showed that the difficulty level of both semantic and non-semantic tasks can be decoded in semantic and domain-general control networks. However, only in the domain-general network, activity patterns generalized across tasks. Gao et al. (2022a) manipulated the association strength between two words (e.g. tea – mug) and found that context-free meaning (word 1) was coded in left aIPL, whereas context-dependent meaning (word 2) for related pairs was represented in left IFG, MFG, pIPL and ventral temporal cortex. Xu et al. (2018) showed that left ATL and temporo-parietal cortex encode taxonomic relationships (e.g. doctor – teacher) and thematic relationships (e.g. doctor – stethoscope). However, in both regions, taxonomic effects were stronger in a taxonomic than thematic task, and vice versa for thematic effects, in support of the view that conceptual representations are enhanced when task-relevant. In summary, previous MVPA studies largely focused on relationships between different concepts, showing that these similarity structures can be modulated by the task or context. Our study extends these previous findings by demonstrating task-dependent modulations of individual perceptual-motor features of concepts (e.g. sound, action features).

Overall, our findings support theories that assume conceptual processing to rely on a flexible, multi-level architecture (Binder and Desai, 2011; Fernandino et al., 2016a; Kemmerer, 2015; Kiefer and Harpaintner, 2020; Popp et al., 2019b). For instance, we previously proposed that conceptual processing relies on a representational hierarchy from modality-specific perceptual-motor regions to multiple levels of cross-modal convergence zones (Kuhnke et al., 2023, 2021, 2020b). The representation of a concept within this neural hierarchy is not a static, task-independent entity, but it is flexibly shaped to the requirements of the current task or context (Hoenig et al., 2008; Kiefer and Pulvermüller, 2012; Lebois et al., 2015; Yee and Thompson-Schill, 2016). Crucially, our current results indicate that the task dependency of conceptual representations varies between different levels of the neural hierarchy: Conceptual representations in modality-specific perceptual-motor regions seem to be selectively retrieved when they are task-relevant (Binder and Desai, 2011; Kuhnke et al., 2021, 2020b). In contrast, conceptual representations in multimodal regions can also be activated (to some extent) when they are task-irrelevant, at least in explicit conceptual judgment tasks (Fernandino et al., 2016a, 2016b).

### 4.2. Cross-modal conceptual representations in multimodal regions

We found that multimodal regions in the bilateral IPL/IPS, left precuneus and left dmPFC allowed for cross-decoding of task-relevant conceptual features: from task-relevant sound to action features, and vice versa. This suggests that these multimodal cortices indeed contain cross-modal representations of task-relevant conceptual information.

Importantly, our results indicate that these cross-modal representations are not “amodal” (i.e., completely invariant to modality-specific features) but “multimodal”, that is, they retain modality-specific information (Kuhnke et al., 2023, 2022, 2020b; see also Fernandino et al., 2022, 2016a, 2016b; Reilly et al., 2016b; Tong et al., 2022). Multimodal regions encode action features (high vs. low action words) during action judgments, and sound features (high vs. low sound words) during sound judgments. Crucially, however, these task-relevant features are represented in an abstract fashion across modalities, encoding their presence vs. absence (cf. Binder, 2016). This multimodal view is supported by several neuroimaging studies (Fernandino et al., 2022, 2016a; Kuhnke et al., 2022, 2020b; Tong et al., 2022). For example, Fernandino et al. (2016a) showed that neural activity in the bilateral IPL, precuneus and dmPFC correlated with the strength of perceptual-motor associations for all tested modalities (action, sound, shape, color, and motion).

As an alternative explanation, cross-decoding between sound and action features could reflect the concomitant activation of action features during sound feature processing, and vice versa (Reilly et al., 2016a). Such “cross-modality spreading” is plausible due to “action–sound coupling”—the phenomenon that many human actions are associated with typical sounds (Lemaitre et al., 2018). However, cross-modality spreading is highly unlikely to explain the cross-modal representations identified in our study. If cross-modality spreading was prevalent, we would have expected cross-modal effects in modality-specific perceptual-motor regions. For example, auditory cortex should have been engaged for action feature retrieval, and somatomotor cortex for sound feature retrieval (Lemaitre et al., 2018; Reilly et al., 2016a). This was clearly not the case: Cross-modal representations were exclusively found in high-level multimodal hubs distant from modality-specific cortices (Binder and Fernandino, 2015; Margulies et al., 2016).

Finally, it should be noted that evidence for a causal role of multimodal conceptual areas is currently weak. For example, we previously found that transcranial magnetic stimulation (TMS) over left IPL selectively impairs action judgments, but not sound judgments, on written words (Kuhnke et al., 2020a). These findings suggest that left IPL might be specialized for action knowledge retrieval, challenging the view of left IPL as a multimodal conceptual hub (also see Ishibashi et al., 2011; Pobric et al., 2010). Future studies should further test the causal relevance of presumptive multimodal regions for the processing of multiple conceptual features.

### 4.3. Involvement of large-scale functional networks in conceptual processing

Our results indicate that conceptual processing mainly recruits the large-scale networks of the default mode network (DMN), frontoparietal control network (FPN) and dorsal attention network (DAN). The searchlight MVPA maps for action and sound feature retrieval, as well as cross-modal representations showed extensive spatial overlap with the DMN, FPN and DAN. In direct network-based decoding analyses, the DMN, FPN and DAN yielded the highest decoding accuracies for task-relevant conceptual features, constituted the only networks that enabled decoding of task-irrelevant features, and showed evidence for cross-modal conceptual representations.

These findings partially support views suggesting a correspondence between the conceptual system—particularly cross-modal convergence zones—and the DMN (Binder et al., 2009, 1999; Fernandino et al., 2016a). However, our results indicate that the DMN is not the only large-scale network engaged in conceptual cognition; conceptual processing also recruits domain-general executive control (FPN) and attention (DAN) networks. Moreover, our findings suggest that not only the DMN contains cross-modal conceptual representations. While almost all large-scale networks enabled above-chance cross-decoding between task-relevant sound and action features, cross-decoding accuracies were highest in DMN, FPN and DAN. Cross-modal conceptual representations seem to be widely distributed throughout the large-scale networks of the human brain.

Crucially, in all networks including the DMN, FPN and DAN, task-relevant features were associated with higher decoding accuracies than task-irrelevant features. This result further corroborates the task dependency of conceptual feature retrieval (Binder and Desai, 2011; Kemmerer, 2015; Kiefer and Harpaintner, 2020; Kuhnke et al., 2021, 2020b). The task dependency of the FPN and DAN is expected. FPN and DAN strongly overlap with “multiple demand” cortex, which has an established role in cognitive control and flexibility (Assem et al., 2020; Duncan, 2010; Wang et al., 2021). Activation level of these areas positively correlates with cognitive demand across a large variety of tasks (Camilleri et al., 2018; Duncan, 2010; Fedorenko et al., 2013), and their activity patterns can encode task-relevant information (Bracci et al., 2017; Cole et al., 2016; Wang et al., 2021).

The task dependency of the DMN is a more intriguing result. The DMN is traditionally characterized as a “task-negative” network, which is deactivated during attention-demanding tasks as compared to the resting state (Fox et al., 2005; Raichle, 2015). Under the task-negative account, the DMN should not be actively engaged in attention-demanding tasks and should not encode task-relevant information (Wang et al., 2021). Our findings are clearly inconsistent with the task-negative view: The DMN showed significant decoding of sound and action features, with the highest decoding accuracies when these features were task-relevant in explicit, attention-demanding conceptual judgment tasks. In contrast to the task-negative view, our results converge with a growing body of evidence that the DMN is actively engaged in demanding cognitive tasks (Crittenden et al., 2015; Smallwood et al., 2021; Sormaz et al., 2018; Wang et al., 2021). We show that the DMN actively supports task-relevant conceptual feature retrieval. This is in line with the view that DMN deactivation during (non-conceptual) attention-demanding tasks as compared to the “resting state” may indeed *reflect* its involvement in conceptual processing (Binder et al., 2009, 1999; Kuhnke et al., 2022). “Resting” can involve spontaneous thought, autobiographical memory, as well as self-referential and introspective processes (Andrews-Hanna, 2012; Smallwood et al., 2021). Crucially, all of these processes may involve the retrieval of conceptual knowledge (Binder et al., 2009, 1999; Kuhnke et al., 2022). Therefore, the DMN may be “deactivated” during non-conceptual attention-demanding tasks, as compared to rest, since the conceptual processing that occurs during rest is interrupted (Kuhnke et al., 2022; Seghier, 2013). Moreover, the DMN may contribute to conceptual processing via its role in mentalizing (Buckner et al., 2008; Christoff et al., 2016), which could be particularly relevant for the processing of abstract concepts (Kiefer et al., 2022; Ulrich et al., 2022). Self-referential processes may also play a role for concrete concepts, particularly during explicit conceptual tasks that involve simulating oneself as experiential agent (e.g. sound or action judgment; Barsalou, 1999).

Our findings converge with a recent study showing that task goals during conceptual feature matching can be decoded from activity patterns in DMN, FPN and DAN (Wang et al., 2021). However, in that study, task goals were confounded with stimulus differences: On each trial, a goal cue (e.g. “color”) preceded a probe–target pair (e.g. “strawberry” – “cherry”). Moreover, task-irrelevant information could not be decoded. Our study addressed these limitations by directly comparing neural activity during different tasks on the *same* stimuli, allowing us to unambiguously attribute activity differences to task (and not stimulus) differences. Our study therefore extends the previous findings by demonstrating task dependency of conceptual representations in DMN, FPN and DAN, even when the stimuli are identical. In addition, we independently manipulated the relevance of both sound and action features to word meaning, which enabled us to test whether a brain region selectively represents task-relevant, or also task-irrelevant features. We showed that DMN, FPN and DAN represent task-irrelevant conceptual feature information. This could reflect a graded task dependency, where task-irrelevant features can be activated—albeit less strongly than task-relevant features—in explicit conceptual judgment tasks (i.e., sound features during action judgments, and action features during sound judgments). Alternatively, it could reflect the active suppression of task-irrelevant features, which is particularly plausible for the FPN and DAN (Corbetta and Shulman, 2002).

Finally, while DMN, FPN and DAN showed the strongest involvement in conceptual processing out of all large-scale networks, the other networks—including modality-specific perceptual-motor systems (e.g. visual, somatomotor)—also enabled decoding of task-relevant conceptual feature representations. Modality-specific perceptual-motor systems and the DMN are located on opposite sides of the “principal gradient of intrinsic connectivity” of the cerebral cortex (Gao et al., 2022b; Margulies et al., 2016; Wang et al., 2020), with DAN and FPN located in-between (see Figure S4). The principal gradient provides a characterization of the cortical hierarchy, reflecting the sensorimotor-to-association axis of cortical association (Sydnor et al., 2021). Taken together, our results suggest that conceptual processing involves virtually *all* levels of the principal gradient of cortical organization—from modality-specific perceptual-motor systems, via attention and control systems, to the default mode network. These findings support theories that assume conceptual processing to rely on a hierarchical, multi-level neural architecture (Binder and Desai, 2011; Fernandino et al., 2016a; Kiefer and Harpaintner, 2020; Kuhnke et al., 2023, 2021, 2020b; Reilly et al., 2016b). Importantly, our results add further evidence that this hierarchical system is flexible, with different levels being engaged in a task-dependent fashion (Binder and Desai, 2011; Kemmerer, 2015; Kuhnke et al., 2020b).

## 5. Conclusion

In conclusion, we found that (1) conceptual representations in modality-specific perceptual-motor and multimodal brain regions are strongly modulated by the task, (2) conceptual representations in several multimodal regions are indeed cross-modal, and (3) conceptual processing recruits the default mode network (DMN), frontoparietal control network (FPN), and dorsal attention network (DAN). Neural representations in all three of these core networks are enhanced for task-relevant (vs. –irrelevant) conceptual features, and enable cross-decoding between modalities. Overall, these findings suggest that large-scale functional brain networks contribute to conceptual processing in a task-dependent and cross-modal fashion.

## Appendix

**Table A1.** Acronyms for brain regions.

|  |  |
| --- | --- |
| a (prefix) | anterior |
| p (prefix) | posterior |
| d (prefix) | dorsal |
| v (prefix) | ventral |
| AAC | auditory association cortex |
| ATL | anterior temporal lobe |
| dmPFC | dorsomedial prefrontal cortex |
| IFG | inferior frontal gyrus |
| IPL | inferior parietal lobe |
| IPS | intraparietal sulcus |
| LTO | lateral temporal-occipital junction |
| MFG | middle frontal gyrus |
| MTG | middle temporal gyrus |
| PCC | posterior cingulate cortex |
| PreCS | precentral sulcus |

## Statements & Declarations

## Acknowledgments

We thank Annika Tjuka for her tremendous help during data acquisition. We also thank Anke Kummer, Nicole Pampus, and Sylvie Neubert for acquiring participants and assisting the fMRI measurements. Moreover, we thank Toralf Mildner for implementing the dual-echo fMRI sequence. Finally, we are grateful to Marie Beaupain and Maike Herrmann for their assistance in stimulus creation and piloting.

## Author Contributions (CRediT)

**Philipp Kuhnke**: Conceptualization, Investigation, Data curation, Formal analysis, Methodology, Visualization, Writing—original draft, Writing—review and editing.

**Markus Kiefer**: Conceptualization, Writing—review and editing.

**Gesa Hartwigsen**: Conceptualization, Funding acquisition, Supervision, Project administration, Writing—review and editing.

