## Supplementary Material for "Conceptual representations in the default, control and attention networks are task-dependent and cross-modal"

(Kuhnke, Kiefer & Hartwigsen 2023 *Brain & Language*)

#### Supplementary Materials and Methods

**Table S1.** Psycholinguistic measures for the four word categories (means; SD in parentheses).

| | **low sound** | **high sound** | **p** | **low action** | **high action** | **p** |
|---|---|---|---|---|---|---|
| **Sound rating** | 1.33 (0.29) | 4.92 (0.62) | **$<10^{-113}$** | 2.92 (1.67) | 3.34 (2.03) | 0.12 |
| **Action rating** | 3.19 (1.64) | 3.4 (1.64) | 0.37 | 1.73 (0.49) | 4.86 (0.45) | **$<10^{-103}$** |
| **Visual rating** | 4.21 (0.54) | 4.07 (0.81) | 0.15 | 4.16 (0.82) | 4.13 (0.52) | 0.77 |
| **Familiarity rating** | 5.52 (0.39) | 5.47 (0.41) | 0.41 | 5.45 (0.42) | 5.55 (0.37) | 0.08 |
| **Letters** | 6.23 (1.61) | 6.36 (2.0) | 0.61 | 6.27 (1.98) | 6.32 (1.63) | 0.84 |
| **Syllables** | 2.21 (0.77) | 2.29 (0.71) | 0.44 | 2.22 (0.71) | 2.28 (0.76) | 0.56 |
| **Lemma freq.** | 5.33 (6.33) | 5.56 (10.35) | 0.85 | 4.74 (6.34) | 6.14 (10.3) | 0.26 |
| **Bigram freq.** | 254220.81 (136859.15) | 231922.15 (122883.01) | 0.24 | 254413.24 (139511.49) | 231729.73 (119826.78) | 0.23 |
| **Trigram freq.** | 145830.18 (85313.42) | 132754.69 (76696.95) | 0.27 | 146068.82 (84690.3) | 132516.05 (77342.95) | 0.25 |
| **Orthographic neighbors** | 7.41 (6.7) | 6.09 (5.86) | 0.15 | 7.05 (6.83) | 6.44 (5.77) | 0.5 |

Ratings were obtained from a total of 163 subjects who did not participate in the fMRI experiment. All other psycholinguistic measures were extracted from the *dlexDB* database (Heister et al., 2011; http://dlexdb.de/). Lemma, bigram and trigram frequencies and number of orthographic neighbors are given per one million words. Freq = frequency.

### Supplementary Results

#### Whole-brain results for task-irrelevant features

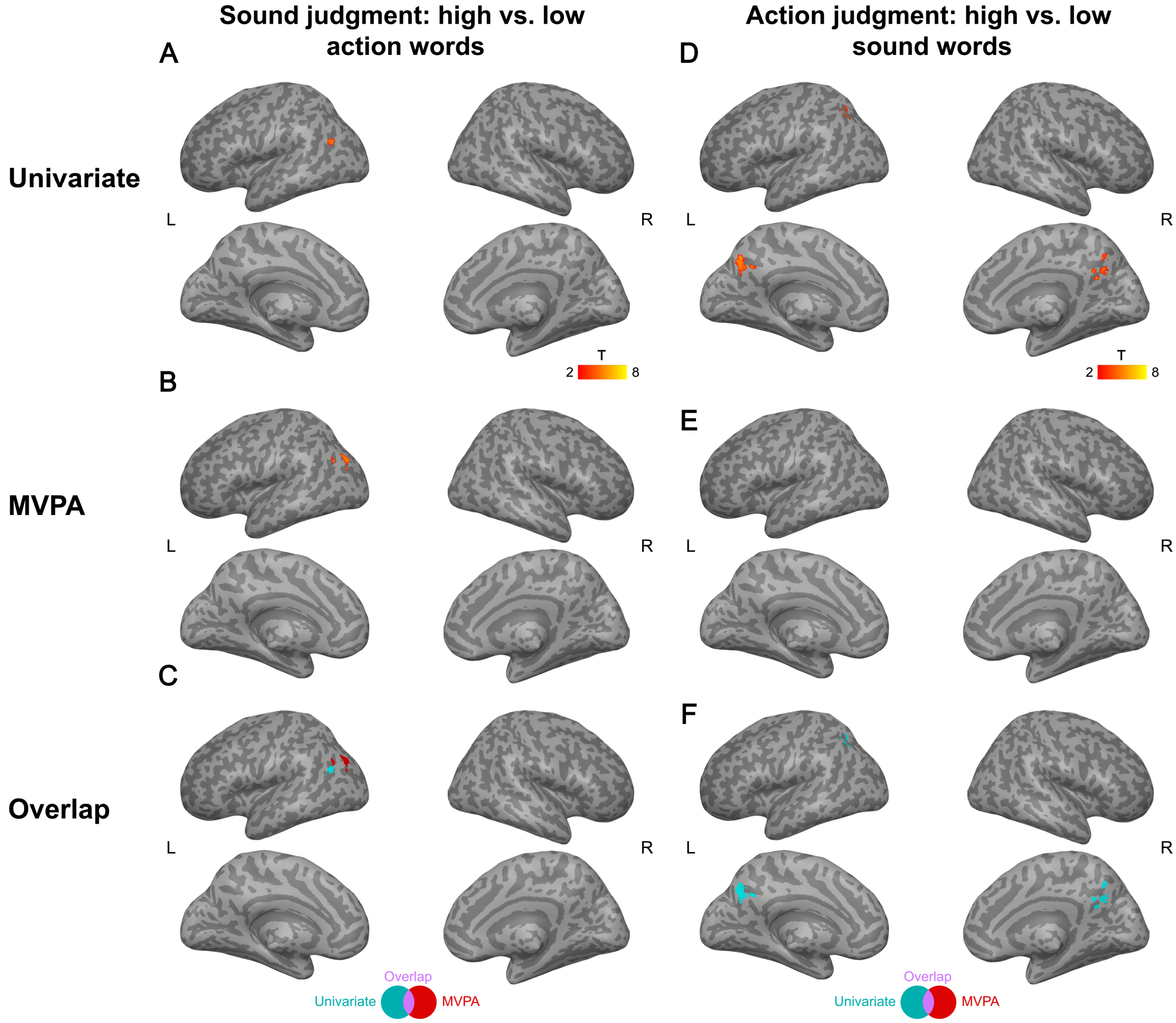


**Figure S1. Comparison of results for whole-brain univariate analysis vs. searchlight MVPA on task-irrelevant conceptual feature retrieval.** Both univariate and MVPA subject-specific maps were smoothed with a 5-mm FWHM Gaussian kernel. All group-level maps were thresholded at a voxel-wise $p < 0.001$ and a cluster-wise $p < 0.05$ FWE-corrected (using non-parametric permutation tests).

#### Supplementary MVPA decoding analyses using 17-network parcellation

In addition to our main MVPA decoding analysis using the 7-network parcellation by Yeo et al. (2011), we also performed each decoding analysis using the more fine-grained 17-network parcellation.

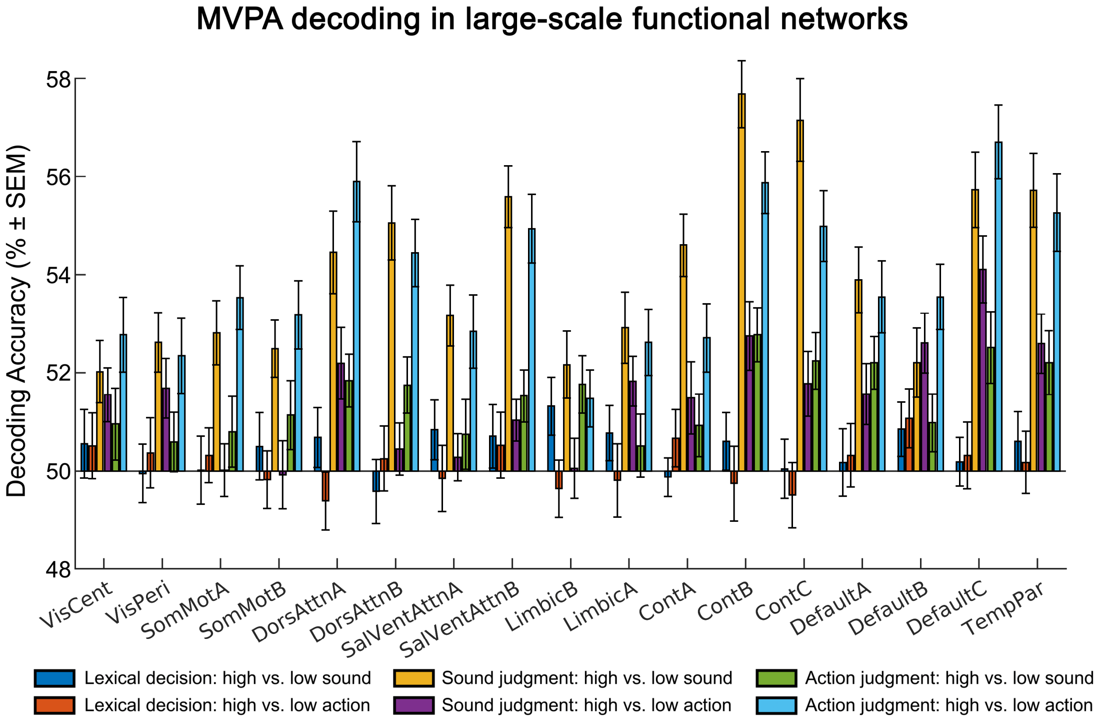


**Figure S2. Results of ROI-based MVPA decoding analyses in the 17 resting-state networks by Yeo et al. (2011).** A machine-learning classifier was trained on the activation patterns in a given network for 5 out of the 6 blocks, and tested on the remaining block (i.e., leave-one-block-out cross validation).

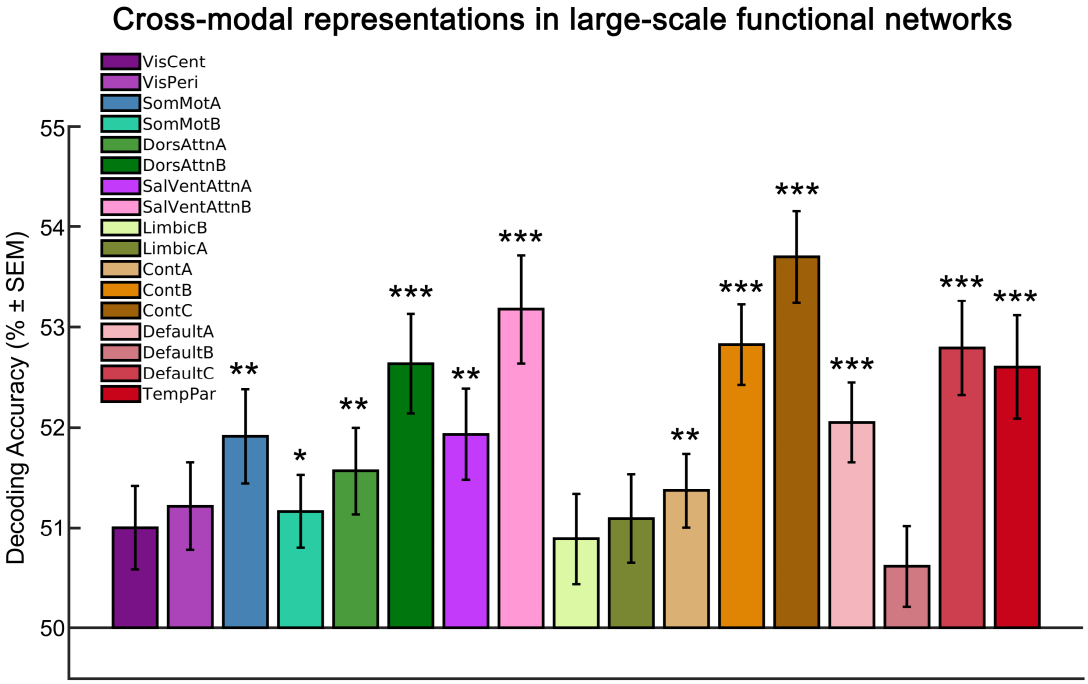


**Figure S3. Cross-decoding of task-relevant conceptual features in the 17 resting-state networks by Yeo et al. (2011).** The classifier was trained on activation patterns for task-relevant sound features (sound judgments: high vs. low sound words) and tested on task-relevant action features (action judgments: high vs. low action words), and vice versa. ***: $p < 0.001$; **: $p < 0.01$; *: $p < 0.05$ (Bonferroni-corrected for the number of networks).

#### Relationship between large-scale networks and the principal gradient of connectivity

As a supplementary analysis, we assessed the relationship between the 7 large-scale networks (Yeo et al., 2011) and the principal gradient of intrinsic connectivity (Gao et al., 2022; Margulies et al., 2016). The principal gradient captures the gradual change of intrinsic connectivity across the cortex (Wang et al., 2020). For each large-scale network, we extracted the mean gradient value across all voxels in the gradient map by Margulies et al. (2016) (available at https://identifiers.org/neurovault.collection:1598) that fell into the respective network ROI.

The results show that the principal gradient extends from modality-specific perceptual-motor systems (visual, somatomotor) on the lower end to the default mode network (DMN) at the upper end, as reported previously (Gao et al., 2022; Margulies et al., 2016; Wang et al., 2020). Dorsal and ventral attention networks, and the limbic and frontoparietal control systems are located at the middle of the gradient.

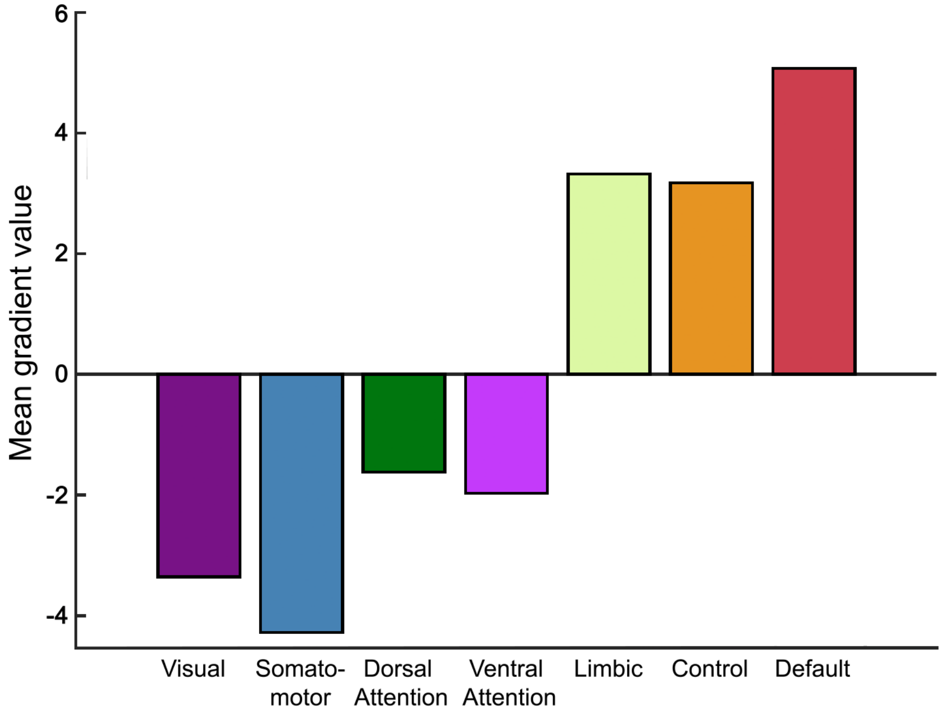


**Figure S4. Gradient values of each large-scale functional network.** The mean gradient value was extracted across all voxels in the gradient map by Margulies et al. (2016) that fell into each large-scale network of the 7-network parcellation by Yeo et al. (2011).

#### Coordinate Tables

The following tables report significant clusters and peak coordinates for the whole-brain univariate and MVPA searchlight analyses. All analyses were thresholded at a voxel-wise $p < 0.001$ and a cluster-wise $p < 0.05$ FWE-corrected using non-parametric permutation tests (5000 permutations). We report up to 3 peaks more than 8 mm apart in clusters larger than 100 $mm^3$. Anatomical labels were determined using the SPM Anatomy toolbox (Version 2.2c; Eickhoff et al., 2005), the Harvard-Oxford atlas distributed with FSL (http://www.fmrib.ox.ac.uk/fsl/), and the human motor area template (http://lrnlab.org/; Mayka et al., 2006). Coordinates are in MNI space.

| | | | |
|---|---|---|---|
| L | left | MFG | middle frontal gyrus |
| R | right | MTG | middle temporal gyrus |
| a (prefix) | anterior | OFC | orbitofrontal cortex |
| p (prefix) | posterior | PCC | posterior cingulate cortex |
| ACC | anterior cingulate cortex | PFC | prefrontal cortex |
| FG | fusiform gyrus | mPFC | medial PFC |
| IFG | inferior frontal gyrus | dmPFC | dorsomedial PFC |
| IFG op | IFG pars opercularis | vmPFC | ventromedial PFC |
| IFG orb | IFG pars orbitalis | PMC | premotor cortex |
| IFG tri | IFG pars triangularis | PMd | dorsal PMC |
| IPL | inferior parietal lobe | PMv | ventral PMC |
| IPS | intraparietal sulcus | PreCS | precentral sulcus |
| ITG | Inferior temporal gyrus | SMA | supplementary motor area |
| LTO | lateral temporal-occipital junction | SPL | superior parietal lobe |
| MCC | middle cingulate cortex | STG | superior temporal gyrus |

**Table S2.** Univariate results for action feature retrieval (action judgments: high vs. low action words).

| Region | Cluster size ($mm^3$) | x | y | z | T |
|---|---|---|---|---|---|
| L IPL/IPS | 12406 | | | | |
| L IPS (hIP2) | | -47 | -44 | 48 | 8.03 |
| L IPL | | -47 | -47 | 58 | 5.81 |
| L IPL (PFm) | | -44 | -54 | 52 | 5.78 |
| L pMTG/ITG | 9750 | | | | |
| L pITG | | -60 | -42 | -18 | 6.65 |
| L pMTG | | -50 | -62 | 2 | 5.40 |
| L pMTG | | -57 | -54 | 2 | 5.37 |
| L aIFG / OFC | 3531 | | | | |
| L OFC | | -47 | 48 | -2 | 5.31 |
| L IFGorb | | -42 | 46 | -15 | 5.08 |
| L OFC | | -42 | 56 | -10 | 5.07 |
| R Cerebellum | 3328 | | | | |
| R Cerebellum (Crus 2) | | 30 | -80 | -45 | 4.58 |
| R Cerebellum (Crus 1) | | 40 | -62 | -40 | 4.47 |
| R Cerebellum (Crus 2) | | 28 | -82 | -42 | 4.40 |
| L aIFG / OFC | 1766 | | | | |
| L IFG orb (Fo3) | | -17 | 23 | -20 | 5.51 |
| L IFG orb (Fo3) | | -22 | 26 | -25 | 4.63 |
| L OFC (Fo2) | | -17 | 18 | -18 | 4.62 |
| L PCC / MCC | 1547 | | | | |
| L PCC | | -2 | -34 | 30 | 5.21 |

| | | | | | |
|---|---|---|---|---|---|
| L MCC | | 0 | -30 | 32 | 5.06 |
| L MCC | | -2 | -27 | 35 | 5.06 |
| L Caudate | 1234 | -10 | 13 | 0 | 6.13 |

**Table S3.** MVPA results for action feature retrieval (action judgments: high vs. low action words).

| Region | Cluster size (mm³) | x | y | z | T |
|---|---|---|---|---|---|
| L IPL/IPS, SPL, pMTG/LTO | 46641 | | | | |
| L IPS (hIP2) | | -47 | -44 | 48 | 6.26 |
| L FG (FG4) | | -42 | -47 | -18 | 6.12 |
| L IPL (PGp) | | -50 | -67 | 30 | 6.10 |
| L IFG, PMC, pre-SMA | 18281 | | | | |
| L IFG tri | | -44 | 43 | 2 | 5.74 |
| L PMv | | -50 | 6 | 40 | 5.70 |
| L IFG tri (44) | | -54 | 16 | 2 | 5.50 |
| L dmPFC, ACC | 8594 | | | | |
| L ACC | | -2 | 48 | 12 | 5.57 |
| L dmPFC (medial SFG) | | -2 | 43 | 35 | 5.17 |
| L ACC | | -12 | 26 | 30 | 5.07 |
| R aIFG | 7047 | | | | |
| R IFG op (45) | | 58 | 18 | 15 | 5.59 |
| R IFG tri | | 40 | 26 | 22 | 5.27 |
| R IFG tri (45) | | 53 | 18 | 22 | 5.04 |
| R Cerebellum | 3094 | | | | |
| R Cerebellum (Crus I) | | 36 | -64 | -40 | 5.79 |
| R Cerebellum (Crus I) | | 40 | -70 | -45 | 5.26 |
| R Cerebellum (Crus II) | | 43 | -72 | -48 | 4.98 |
| R MFG | 2609 | | | | |
| R MFG | | 28 | 40 | 28 | 4.77 |
| R MFG | | 33 | 40 | 25 | 4.72 |
| R MFG | | 33 | 50 | 28 | 4.40 |
| L PCC / Precuneus | 1906 | | | | |
| L PCC | | -2 | -34 | 35 | 4.78 |
| L Precuneus | | -7 | -54 | 45 | 4.44 |
| L Precuneus | | -4 | -50 | 40 | 4.25 |
| L Cerebellum | 1844 | | | | |
| L Cerebellum (Crus II) | | -40 | -77 | -50 | 4.56 |
| L Cerebellum (lobule VIIIa) | | -10 | -74 | -52 | 4.36 |
| L Cerebellum (lobule VIIb) | | -32 | -67 | -50 | 4.04 |
| R Calcarine Gyrus | 1813 | | | | |
| R Calcarine Gyrus | | 6 | -57 | 15 | 4.99 |
| R Calcarine Gyrus (hOc2) | | 16 | -52 | 8 | 4.43 |
| R Calcarine Gyrus (hOc2) | | -2 | -64 | 12 | 3.99 |
| L/R Cerebellum | 1547 | | | | |
| Cerebellar Vermis (lobule VIIIa) | | 0 | -60 | -30 | 5.33 |
| Cerebellar Vermis (lobule VIIIa) | | 0 | -67 | -30 | 4.45 |
| R Cerebellum (lobule VI) | | 8 | -77 | -28 | 3.93 |
| R IPL/IPS, SPL, S1 | 1531 | | | | |
| R IPS (hIP3) | | 38 | -54 | 52 | 5.02 |
| R IPL (PGa) | | 50 | -60 | 40 | 4.05 |
| R SPL (7PC) | | 36 | -47 | 58 | 3.86 |

| | | | | | |
|---|---|---|---|---|---|
| R IPL | 1453 | | | | |
| R IPL (PGp) | | 46 | -67 | 30 | 4.41 |
| R IPL (PGp) | | 48 | -82 | 20 | 3.72 |
| R IPL (PGp) | | 58 | -67 | 32 | 3.71 |
| R MFG | 1344 | | | | |
| R MFG | | 30 | 16 | 42 | 5.44 |
| R MFG | | 28 | 28 | 45 | 4.82 |
| R MFG | | 33 | 20 | 48 | 4.01 |

**Table S4.** Overlap between univariate and MVPA results for action feature retrieval (action judgments: high vs. low action words).

| Region | Cluster size (mm$^3$) | x | y | z | T |
|---|---|---|---|---|---|
| L IPL/IPS, SPL | 7813 | | | | |
| L IPS (hIP2) | | -47 | -44 | 48 | 6.26 |
| L IPL (PFm) | | -42 | -60 | 50 | 5.30 |
| L IPL (PFt) | | -60 | -34 | 42 | 4.15 |
| L pMTG | 1828 | | | | |
| L pMTG | | -50 | -60 | 2 | 4.96 |
| L pMTG | | -60 | -47 | 2 | 4.09 |
| L pMTG | | -64 | -50 | -5 | 4.02 |
| R Cerebellum | 1266 | | | | |
| R Cerebellum (Crus I) | | 38 | -62 | -40 | 4.47 |
| R Cerebellum (Crus I) | | 38 | -70 | -42 | 4.27 |
| R Cerebellum (Crus I) | | 46 | -67 | -40 | 3.99 |
| L pITG / FG | 1156 | | | | |
| L pITG | | -57 | -40 | -15 | 4.95 |
| L FG (FG4) | | -50 | -47 | -18 | 3.85 |
| L aIFG / OFC, MFG | 500 | | | | |
| L OFC | | -47 | 46 | -2 | 4.09 |
| L MFG | | -44 | 50 | 2 | 3.79 |
| L IFG tri | | -44 | 46 | 5 | 3.58 |
| L PCC | 438 | -2 | -32 | 32 | 4.49 |
| L FG (FG4) | 47 | -50 | -60 | -12 | 3.77 |

**Table S5.** Univariate results for sound feature retrieval (sound judgments: high vs. low sound words).

| Region | Cluster size (mm$^3$) | x | y | z | T |
|---|---|---|---|---|---|
| L IPL/IPS | 9438 | | | | |
| L IPL (PGa) | | -44 | -62 | 50 | 5.96 |
| L IPL (PGa) | | -37 | -72 | 50 | 5.23 |
| L IPS (hIP2) | | -50 | -44 | 52 | 5.11 |
| L aIFG | 8250 | | | | |
| L IFG orb | | -47 | 46 | -8 | 5.68 |
| L IFG orb | | -37 | 36 | -18 | 5.31 |
| L IFG tri | | -47 | 46 | 2 | 4.87 |
| L dmPFC | 7188 | | | | |
| L dmPFC (medial SFG) | | -4 | 33 | 38 | 6.94 |
| L dmPFC (medial SFG) | | -12 | 48 | 38 | 5.53 |
| L dmPFC (medial SFG) | | -4 | 30 | 48 | 5.49 |
| L MFG / PreCS | 3078 | | | | |
| L MFG | | -47 | 16 | 42 | 5.17 |
| L MFG | | -40 | 10 | 48 | 4.80 |
| L PreCS | | -47 | 20 | 30 | 4.54 |
| L vmPFC | 1297 | | | | |
| L vmPFC (Fo2) | | -12 | 18 | -18 | 5.92 |
| L vmPFC (Fo3) | | -14 | 23 | -22 | 4.58 |
| L vmPFC (Fo3) | | -17 | 33 | -22 | 4.57 |
| L pMTG | 1141 | | | | |
| L pMTG | | -57 | -42 | -12 | 5.26 |

| | | | | | |
|---|---|---|---|---|---|
| L pMTG | | -62 | -40 | -2 | 4.15 |
| L pMTG | | -60 | -44 | -5 | 3.59 |
| R Cerebellum | 969 | | | | |
| R Cerebellum (Crus I) | | 40 | -64 | -42 | 4.29 |
| R Cerebellum (Crus II) | | 40 | -74 | -45 | 4.06 |
| R Cerebellum (Crus II) | | 36 | -77 | -45 | 4.01 |
| L Insula | 813 | | | | |
| L Insula | | -30 | 23 | -10 | 5.29 |
| L Insula | | -27 | 26 | -5 | 4.35 |
| L Insula | | -30 | 18 | 5 | 4.08 |

**Table S6.** MVPA results for sound feature retrieval (sound judgments: high vs. low sound words).

| Region | Cluster size ($mm^3$) | x | y | z | T |
|---|---|---|---|---|---|
| L/R mPFC / ACC, aIFG | 83406 | | | | |
| L dmPFC (medial SFG) | | -20 | 36 | 55 | 7.22 |
| L dmPFC | | -24 | 16 | 58 | 6.56 |
| L dmPFC (medial SFG) | | -12 | 46 | 40 | 6.27 |
| L/R IPL/IPS, Precuneus / PCC | 61391 | | | | |
| L IPS (hIP2) | | -44 | -50 | 45 | 8.34 |
| L IPL (PFt) | | -52 | -37 | 42 | 7.33 |
| L IPS (hIP3) | | -30 | -62 | 50 | 6.86 |
| R IPL/IPS, SPL | 10219 | | | | |
| R IPL (PF) | | 68 | -24 | 38 | 6.36 |
| R IPL (PFm) | | 48 | -54 | 55 | 6.15 |
| R IPL (PFm) | | 50 | -44 | 38 | 5.88 |
| L AAC (pSTG/MTG), FG | 9031 | | | | |
| L MOG (hOc5) | | -50 | -67 | -2 | 5.92 |
| L FG (FG4) | | -50 | -54 | -12 | 5.50 |
| L FG (FG4) | | -47 | -50 | -25 | 5.48 |
| R aIFG, Insula | 5797 | | | | |
| R IFG tri (45) | | 56 | 40 | 5 | 6.97 |
| R Insula | | 33 | 28 | 0 | 5.33 |
| R IFG tri (45) | | 48 | 38 | 15 | 5.08 |
| R AAC (pSTG/MTG), IPL | 2734 | | | | |
| R AAC (TE3) | | 58 | -37 | 2 | 5.75 |
| R AAC (TE3) | | 68 | -27 | 2 | 5.59 |
| R IPL (PF) | | 60 | -40 | 15 | 3.68 |

**Table S7.** Overlap between univariate and MVPA results for sound feature retrieval (sound judgments: high vs. low sound words).

| Region | Cluster size ($mm^3$) | x | y | z | T |
|---|---|---|---|---|---|
| L IPL/IPS | 8172 | | | | |
| L IPL (PFm) | | -42 | -60 | 50 | 5.21 |
| L IPS (hIP2) | | -47 | -44 | 50 | 5.01 |
| L IPS (hIP3) | | -34 | -57 | 48 | 4.98 |
| L dmPFC | 5375 | | | | |
| L dmPFC (medial SFG) | | -12 | 48 | 38 | 5.37 |
| L dmPFC (medial SFG) | | -2 | 36 | 35 | 5.23 |
| L dmPFC (medial SFG) | | -4 | 28 | 38 | 5.16 |
| L aIFG | 4609 | | | | |
| L IFG orb | | -42 | 48 | -10 | 4.60 |
| L MFG | | -44 | 50 | 8 | 4.59 |
| L IFG orb | | -42 | 53 | -2 | 4.51 |
| L MFG / PreCS | 1844 | | | | |
| L MFG | | -47 | 13 | 42 | 4.59 |
| L MFG | | -37 | 10 | 50 | 4.12 |
| L PreCS | | -50 | 18 | 32 | 4.09 |
| L pMTG | 859 | | | | |
| L pMTG | | -60 | -42 | -12 | 4.68 |
| L pMTG | | -60 | -44 | -5 | 3.58 |

| | | | | | |
|---|---|---|---|---|---|
| L Insula | 578 | | | | |
| L Insula | | -30 | 23 | -8 | 4.36 |
| L Insula | | -32 | 20 | -10 | 4.23 |
| L dmPFC | 125 | -2 | 16 | 60 | 4.22 |
| L IFG orb (Fo3) | 109 | -34 | 38 | -18 | 3.67 |

**Table S8.** Univariate results for multimodal regions (conjunction of [action judgments: high vs. low action words] ∩ [sound judgments: high vs. low sound words]).

| **Region** | **Cluster size (mm$^3$)** | **x** | **y** | **z** | **T** |
|---|---|---|---|---|---|
| L IPL/IPS | 4188 | | | | |
| L IPL (PFm) | | -42 | -60 | 50 | 5.30 |
| L IPS (hIP2) | | -50 | -44 | 52 | 5.11 |
| L IPS (hIP3) | | -40 | -62 | 42 | 4.16 |
| L aIFG | 2266 | | | | |
| L IFG orb | | -47 | 48 | -2 | 4.78 |
| L IFG orb | | -44 | 46 | -15 | 4.77 |
| L IFG orb | | -37 | 46 | -15 | 4.45 |
| L pMTG/ITG | 906 | | | | |
| L pITG | | -57 | -42 | -15 | 4.87 |
| L pMTG | | -62 | -44 | -5 | 3.55 |
| R Cerebellum | 703 | | | | |
| R Cerebellum (Crus II) | | 36 | -77 | -45 | 4.01 |
| R Cerebellum (Crus I) | | 40 | -67 | -42 | 3.91 |
| L vmPFC | 375 | | | | |
| L vmPFC (Fo2) | | -14 | 20 | -20 | 4.24 |
| L vmPFC (Fo3) | | -20 | 28 | -20 | 3.88 |
| L vmPFC (Fo3) | | -17 | 26 | -22 | 3.69 |

**Table S9.** MVPA results for multimodal regions (conjunction of [action judgments: high vs. low action words] ∩ [sound judgments: high vs. low sound words]).

| **Region** | **Cluster size (mm$^3$)** | **x** | **y** | **z** | **T** |
|---|---|---|---|---|---|
| L IPL/IPS, Precuneus | 18422 | | | | |
| L IPS (hIP2) | | -47 | -44 | 48 | 6.17 |
| L IPS (hIP1) | | -42 | -54 | 45 | 5.52 |
| L Precuneus (7A) | | -27 | -70 | 42 | 4.70 |
| L IFG | 6469 | | | | |
| L IFG tri | | -47 | 40 | 5 | 4.70 |
| L IFG op (44) | | -54 | 13 | 2 | 4.69 |
| L IFG tri (45) | | -50 | 43 | 2 | 4.68 |
| L MFG / PreCS | 5234 | | | | |
| L PreCS | | -50 | 10 | 40 | 4.48 |
| L PreCS | | -44 | 3 | 38 | 4.11 |
| L PreCS | | -47 | 3 | 52 | 4.07 |
| L dmPFC / ACC | 2703 | | | | |
| L dmPFC (medial SFG) | | -2 | 40 | 35 | 4.83 |
| L ACC | | -4 | 26 | 32 | 4.80 |
| L ACC | | -12 | 30 | 28 | 4.23 |
| L pMTG/ITG, FG | 2359 | | | | |
| L FG (FG4) | | -42 | -50 | -25 | 4.65 |
| L pMTG | | -64 | -52 | -2 | 4.37 |
| L FG (FG4) | | -44 | -47 | -22 | 4.36 |
| L PCC | 891 | | | | |
| L PCC | | -2 | -34 | 35 | 4.50 |
| L PCC | | -2 | -30 | 35 | 4.44 |
| L PCC | | -7 | -44 | 38 | 3.35 |
| L pITG | 875 | -57 | -40 | -15 | 4.95 |
| L dmPFC / ACC | 688 | | | | |
| L ACC | | -4 | 48 | 12 | 4.42 |
| L ACC | | -2 | 46 | 10 | 4.30 |
| L dmPFC (medial SFG) | | -7 | 43 | 18 | 3.72 |

| | | | | | |
|---|---|---|---|---|---|
| R IPL/IPS | 609 | | | | |
| R IPS (hIP3) | | 40 | -54 | 52 | 4.67 |
| R IPL (PGa) | | 48 | -57 | 45 | 3.46 |
| R aIFG | 531 | | | | |
| R IFG tri (45) | | 48 | 36 | 15 | 4.15 |
| R IFG tri (45) | | 50 | 38 | 12 | 3.95 |
| R IFG tri (45) | | 40 | 30 | 15 | 3.91 |
| R MFG / SFG | 469 | | | | |
| R MFG | | 28 | 20 | 45 | 3.97 |
| R SFG | | 28 | 16 | 38 | 3.91 |
| R MFG | | 28 | 26 | 45 | 3.84 |
| L dmPFC (medial SFG) | 266 | -4 | 56 | 35 | 4.14 |
| L Precuneus | 156 | | | | |
| L Precuneus | | -2 | -50 | 40 | 3.79 |
| L Precuneus | | -7 | -52 | 45 | 3.72 |
| L mPFC (Fp1) | 156 | -7 | 63 | 22 | 3.69 |

**Table S10.** Overlap between univariate and MVPA results for multimodal regions (conjunction of [action judgments: high vs. low action words] ∩ [sound judgments: high vs. low sound words]).

| **Region** | **Cluster size (mm$^3$)** | **x** | **y** | **z** | **T** |
|---|---|---|---|---|---|
| L IPL/IPS | 3453 | | | | |
| L IPL (PFm) | | -42 | -60 | 50 | 5.21 |
| L IPS (hIP2) | | -47 | -44 | 50 | 5.01 |
| L IPS (hIP1) | | -44 | -54 | 48 | 4.67 |
| L pMTG | 453 | -57 | -40 | -12 | 4.66 |
| L aIFG | 344 | | | | |
| L IFG orb | | -44 | 50 | 2 | 3.79 |
| L IFG tri (45) | | -50 | 46 | 0 | 3.72 |
| L IFG tri | | -44 | 46 | 5 | 3.58 |

**Table S11.** Cross-decoding between sound and action feature retrieval (training on [action judgments: high vs. low action words], and testing on [sound judgments: high vs. low sound words], and vice versa).

| **Region** | **Cluster size (mm$^3$)** | **x** | **y** | **z** | **T** |
|---|---|---|---|---|---|
| L IPL/IPS, SPL/S1 | 11094 | | | | |
| L SPL (7A) | | -34 | -54 | 52 | 6.28 |
| L S1 (area 2) | | -42 | -37 | 52 | 5.79 |
| L IPL (PFm) | | -47 | -54 | 55 | 5.30 |
| L Precuneus | 4359 | | | | |
| L Precuneus (7P) | | -10 | -74 | 50 | 6.13 |
| L Precuneus (7A) | | -4 | -70 | 30 | 4.43 |
| L Precuneus | | 0 | -60 | 38 | 4.03 |
| L dmPFC | 3641 | | | | |
| L dmPFC (medial SFG) | | -4 | 33 | 32 | 5.17 |
| L dmPFC (medial SFG) | | 0 | 33 | 42 | 4.63 |
| L dmPFC (medial SFG) | | -10 | 28 | 50 | 4.42 |
| R IPL/IPS, SPL/S1 | 2766 | | | | |
| R S1 (area 2) | | 38 | -40 | 55 | 5.76 |
| R IPL (PFm) | | 50 | -40 | 45 | 4.78 |
| R IPL (PFt) | | 50 | -32 | 38 | 4.32 |
| L aSTG, IFG | 1281 | | | | |
| L IFG (44) | | -52 | 10 | -2 | 4.87 |
| L aSTG | | -47 | 3 | -2 | 3.83 |

#### Comparison of univariate analysis and MVPA decoding in anatomical ROIs

The following tables report the results of statistical analyses on the univariate activation magnitudes and MVPA decoding accuracies in anatomical regions-of-interest (ROIs) commonly implicated in conceptual-semantic processing. For statistical inference, we first conducted one-sample t-tests in each ROI. Second, we conducted paired t-tests comparing the activation magnitudes / decoding accuracies between experimental conditions. P-values were corrected for multiple comparisons using Bonferroni correction for the number of ROIs.

**Table S12.** One-sample t-tests on the univariate activation magnitudes (contrast values in arbitrary units) in each ROI against 0.

| ROI | Analysis | Mean | SD | T | p (corr.) |
|---|---|---|---|---|---|
| **L aIFG** | Lexical decision: high vs. low sound | 0.023 | 0.055 | 0.408 | 1.000 |
| **L aIFG** | Lexical decision: high vs. low action | -0.026 | 0.046 | -0.555 | 1.000 |
| **L aIFG** | Sound judgment: high vs. low sound | 0.247 | 0.069 | 3.572 | **0.003** |
| **L aIFG** | Sound judgment: high vs. low action | -0.225 | 0.052 | -4.307 | 1.000 |
| **L aIFG** | Action judgment: high vs. low sound | -0.031 | 0.057 | -0.553 | 1.000 |
| **L aIFG** | Action judgment: high vs. low action | 0.115 | 0.069 | 1.676 | 0.356 |
| **L MFG** | Lexical decision: high vs. low sound | 0.006 | 0.046 | 0.134 | 1.000 |
| **L MFG** | Lexical decision: high vs. low action | 0.009 | 0.041 | 0.225 | 1.000 |
| **L MFG** | Sound judgment: high vs. low sound | 0.150 | 0.062 | 2.402 | 0.074 |
| **L MFG** | Sound judgment: high vs. low action | 0.012 | 0.058 | 0.211 | 1.000 |
| **L MFG** | Action judgment: high vs. low sound | 0.079 | 0.044 | 1.788 | 0.285 |
| **L MFG** | Action judgment: high vs. low action | 0.091 | 0.052 | 1.766 | 0.299 |
| **L ATL** | Lexical decision: high vs. low sound | 0.041 | 0.048 | 0.846 | 1.000 |
| **L ATL** | Lexical decision: high vs. low action | 0.035 | 0.032 | 1.090 | 0.989 |
| **L ATL** | Sound judgment: high vs. low sound | -0.084 | 0.060 | -1.386 | 1.000 |
| **L ATL** | Sound judgment: high vs. low action | -0.008 | 0.045 | -0.182 | 1.000 |
| **L ATL** | Action judgment: high vs. low sound | 0.068 | 0.051 | 1.331 | 0.668 |
| **L ATL** | Action judgment: high vs. low action | -0.021 | 0.047 | -0.438 | 1.000 |
| **L pMTG** | Lexical decision: high vs. low sound | 0.023 | 0.044 | 0.531 | 1.000 |
| **L pMTG** | Lexical decision: high vs. low action | 0.026 | 0.038 | 0.692 | 1.000 |
| **L pMTG** | Sound judgment: high vs. low sound | 0.057 | 0.051 | 1.106 | 0.964 |
| **L pMTG** | Sound judgment: high vs. low action | -0.012 | 0.045 | -0.256 | 1.000 |
| **L pMTG** | Action judgment: high vs. low sound | 0.110 | 0.042 | 2.617 | **0.044** |
| **L pMTG** | Action judgment: high vs. low action | 0.124 | 0.040 | 3.105 | **0.012** |
| **L pIPL** | Lexical decision: high vs. low sound | 0.035 | 0.044 | 0.785 | 1.000 |
| **L pIPL** | Lexical decision: high vs. low action | 0.037 | 0.039 | 0.958 | 1.000 |
| **L pIPL** | Sound judgment: high vs. low sound | 0.086 | 0.051 | 1.678 | 0.354 |
| **L pIPL** | Sound judgment: high vs. low action | 0.063 | 0.047 | 1.348 | 0.649 |
| **L pIPL** | Action judgment: high vs. low sound | 0.067 | 0.051 | 1.314 | 0.688 |
| **L pIPL** | Action judgment: high vs. low action | 0.197 | 0.046 | 4.271 | **<0.001** |
| **PCC/Precuneus** | Lexical decision: high vs. low sound | 0.009 | 0.057 | 0.150 | 1.000 |
| **PCC/Precuneus** | Lexical decision: high vs. low action | 0.037 | 0.043 | 0.858 | 1.000 |
| **PCC/Precuneus** | Sound judgment: high vs. low sound | -0.016 | 0.078 | -0.212 | 1.000 |
| **PCC/Precuneus** | Sound judgment: high vs. low action | 0.076 | 0.057 | 1.325 | 0.676 |
| **PCC/Precuneus** | Action judgment: high vs. low sound | 0.123 | 0.054 | 2.289 | 0.097 |
| **PCC/Precuneus** | Action judgment: high vs. low action | 0.019 | 0.053 | 0.354 | 1.000 |
| **dmPFC** | Lexical decision: high vs. low sound | 0.003 | 0.038 | 0.086 | 1.000 |
| **dmPFC** | Lexical decision: high vs. low action | 0.014 | 0.032 | 0.436 | 1.000 |
| **dmPFC** | Sound judgment: high vs. low sound | 0.043 | 0.050 | 0.843 | 1.000 |
| **dmPFC** | Sound judgment: high vs. low action | -0.021 | 0.044 | -0.477 | 1.000 |
| **dmPFC** | Action judgment: high vs. low sound | 0.028 | 0.037 | 0.749 | 1.000 |
| **dmPFC** | Action judgment: high vs. low action | 0.034 | 0.035 | 0.961 | 1.000 |

**Table S13.** Paired t-tests comparing the univariate activation magnitudes between conditions within each ROI.

| ROI | Comparison | Mean | SD | T | p (corr.) |
|---|---|---|---|---|---|
| **L aIFG** | Lexical decision vs. action judgment for high vs. low action | -0.141 | 0.520 | -1.715 | 0.660 |
| **L aIFG** | Lexical decision vs. action judgment for high vs. low sound | 0.054 | 0.460 | 0.745 | 1.000 |

| | | | | | |
|---|---|---|---|---|---|
| **L aIFG** | Lexical decision vs. sound judgment for high vs. low action | 0.200 | 0.399 | 3.166 | **0.021** |
| **L aIFG** | Lexical decision vs. sound judgment for high vs. low sound | -0.225 | 0.496 | -2.864 | **0.047** |
| **L aIFG** | Lexical decision: sound vs. action | 0.048 | 0.559 | 0.546 | 1.000 |
| **L aIFG** | Sound judgment vs. action judgment for high vs. low action | -0.341 | 0.416 | -5.181 | **<0.001** |
| **L aIFG** | Sound judgment vs. action judgment for high vs. low sound | 0.279 | 0.605 | 2.914 | **0.041** |
| **L aIFG** | Sound judgment: sound vs. action | 0.473 | 0.625 | 4.781 | **<0.001** |
| **L aIFG** | Action judgment: sound vs. action | -0.147 | 0.552 | -1.682 | 0.703 |
| **L MFG** | Lexical decision vs. action judgment for high vs. low action | -0.082 | 0.412 | -1.261 | 1.000 |
| **L MFG** | Lexical decision vs. action judgment for high vs. low sound | -0.073 | 0.353 | -1.301 | 1.000 |
| **L MFG** | Lexical decision vs. sound judgment for high vs. low action | -0.003 | 0.451 | -0.042 | 1.000 |
| **L MFG** | Lexical decision vs. sound judgment for high vs. low sound | -0.143 | 0.463 | -1.957 | 0.402 |
| **L MFG** | Lexical decision: sound vs. action | -0.003 | 0.459 | -0.042 | 1.000 |
| **L MFG** | Sound judgment vs. action judgment for high vs. low action | -0.079 | 0.340 | -1.471 | 1.000 |
| **L MFG** | Sound judgment vs. action judgment for high vs. low sound | 0.071 | 0.462 | 0.970 | 1.000 |
| **L MFG** | Sound judgment: sound vs. action | 0.137 | 0.584 | 1.489 | 1.000 |
| **L MFG** | Action judgment: sound vs. action | -0.013 | 0.448 | -0.177 | 1.000 |
| **L ATL** | Lexical decision vs. action judgment for high vs. low action | 0.055 | 0.352 | 0.994 | 1.000 |
| **L ATL** | Lexical decision vs. action judgment for high vs. low sound | -0.027 | 0.348 | -0.491 | 1.000 |
| **L ATL** | Lexical decision vs. sound judgment for high vs. low action | 0.043 | 0.371 | 0.735 | 1.000 |
| **L ATL** | Lexical decision vs. sound judgment for high vs. low sound | 0.125 | 0.488 | 1.614 | 0.802 |
| **L ATL** | Lexical decision: sound vs. action | 0.006 | 0.400 | 0.095 | 1.000 |
| **L ATL** | Sound judgment vs. action judgment for high vs. low action | 0.012 | 0.406 | 0.191 | 1.000 |
| **L ATL** | Sound judgment vs. action judgment for high vs. low sound | -0.152 | 0.432 | -2.218 | 0.227 |
| **L ATL** | Sound judgment: sound vs. action | -0.076 | 0.482 | -0.992 | 1.000 |
| **L ATL** | Action judgment: sound vs. action | 0.088 | 0.419 | 1.335 | 1.000 |
| **L pMTG** | Lexical decision vs. action judgment for high vs. low action | -0.098 | 0.361 | -1.715 | 0.660 |
| **L pMTG** | Lexical decision vs. action judgment for high vs. low sound | -0.086 | 0.267 | -2.042 | 0.335 |
| **L pMTG** | Lexical decision vs. sound judgment for high vs. low action | 0.038 | 0.420 | 0.566 | 1.000 |
| **L pMTG** | Lexical decision vs. sound judgment for high vs. low sound | -0.034 | 0.448 | -0.475 | 1.000 |
| **L pMTG** | Lexical decision: sound vs. action | -0.003 | 0.405 | -0.044 | 1.000 |
| **L pMTG** | Sound judgment vs. action judgment for high vs. low action | -0.135 | 0.322 | -2.657 | 0.079 |
| **L pMTG** | Sound judgment vs. action judgment for high vs. low sound | -0.053 | 0.434 | -0.767 | 1.000 |
| **L pMTG** | Sound judgment: sound vs. action | 0.068 | 0.410 | 1.054 | 1.000 |
| **L pMTG** | Action judgment: sound vs. action | -0.014 | 0.394 | -0.232 | 1.000 |
| **L pIPL** | Lexical decision vs. action judgment for high vs. low action | -0.160 | 0.389 | -2.594 | 0.093 |
| **L pIPL** | Lexical decision vs. action judgment for high vs. low sound | -0.033 | 0.388 | -0.536 | 1.000 |
| **L pIPL** | Lexical decision vs. sound judgment for high vs. low action | -0.026 | 0.415 | -0.394 | 1.000 |
| **L pIPL** | Lexical decision vs. sound judgment for high vs. low sound | -0.051 | 0.465 | -0.694 | 1.000 |
| **L pIPL** | Lexical decision: sound vs. action | -0.003 | 0.431 | -0.041 | 1.000 |
| **L pIPL** | Sound judgment vs. action judgment for high vs. low action | -0.134 | 0.364 | -2.324 | 0.178 |
| **L pIPL** | Sound judgment vs. action judgment for high vs. low sound | 0.018 | 0.479 | 0.239 | 1.000 |
| **L pIPL** | Sound judgment: sound vs. action | 0.022 | 0.464 | 0.304 | 1.000 |
| **L pIPL** | Action judgment: sound vs. action | -0.129 | 0.446 | -1.836 | 0.518 |

| | | | | | |
|---|---|---|---|---|---|
| **PCC/Precuneus** | Lexical decision vs. action judgment for high vs. low action | 0.018 | 0.441 | 0.261 | 1.000 |
| **PCC/Precuneus** | Lexical decision vs. action judgment for high vs. low sound | -0.115 | 0.418 | -1.733 | 0.638 |
| **PCC/Precuneus** | Lexical decision vs. sound judgment for high vs. low action | -0.039 | 0.492 | -0.498 | 1.000 |
| **PCC/Precuneus** | Lexical decision vs. sound judgment for high vs. low sound | 0.025 | 0.657 | 0.240 | 1.000 |
| **PCC/Precuneus** | Lexical decision: sound vs. action | -0.028 | 0.499 | -0.360 | 1.000 |
| **PCC/Precuneus** | Sound judgment vs. action judgment for high vs. low action | 0.057 | 0.350 | 1.028 | 1.000 |
| **PCC/Precuneus** | Sound judgment vs. action judgment for high vs. low sound | -0.139 | 0.593 | -1.488 | 1.000 |
| **PCC/Precuneus** | Sound judgment: sound vs. action | -0.092 | 0.619 | -0.941 | 1.000 |
| **PCC/Precuneus** | Action judgment: sound vs. action | 0.104 | 0.496 | 1.330 | 1.000 |
| **dmPFC** | Lexical decision vs. action judgment for high vs. low action | -0.020 | 0.302 | -0.414 | 1.000 |
| **dmPFC** | Lexical decision vs. action judgment for high vs. low sound | -0.024 | 0.294 | -0.523 | 1.000 |
| **dmPFC** | Lexical decision vs. sound judgment for high vs. low action | 0.035 | 0.349 | 0.631 | 1.000 |
| **dmPFC** | Lexical decision vs. sound judgment for high vs. low sound | -0.039 | 0.368 | -0.674 | 1.000 |
| **dmPFC** | Lexical decision: sound vs. action | -0.011 | 0.368 | -0.184 | 1.000 |
| **dmPFC** | Sound judgment vs. action judgment for high vs. low action | -0.055 | 0.257 | -1.345 | 1.000 |
| **dmPFC** | Sound judgment vs. action judgment for high vs. low sound | 0.015 | 0.397 | 0.238 | 1.000 |
| **dmPFC** | Sound judgment: sound vs. action | 0.063 | 0.454 | 0.882 | 1.000 |
| **dmPFC** | Action judgment: sound vs. action | -0.006 | 0.305 | -0.128 | 1.000 |

**Table S14.** One-sample t-tests on the MVPA decoding accuracies in each ROI against chance level (50%).

| ROI | Analysis | Mean | SD | T | p (corr.) |
|---|---|---|---|---|---|
| **L aIFG** | Lexical decision: high vs. low sound | -0.195 | 0.803 | -0.243 | 1.000 |
| **L aIFG** | Lexical decision: high vs. low action | 0.182 | 0.687 | 0.265 | 1.000 |
| **L aIFG** | Sound judgment: high vs. low sound | 3.997 | 0.659 | 6.064 | **<0.001** |
| **L aIFG** | Sound judgment: high vs. low action | 0.169 | 0.672 | 0.252 | 1.000 |
| **L aIFG** | Action judgment: high vs. low sound | 2.044 | 0.6 | 3.407 | **0.005** |
| **L aIFG** | Action judgment: high vs. low action | 3.88 | 0.585 | 6.63 | **<0.001** |
| **L MFG** | Lexical decision: high vs. low sound | -0.182 | 0.599 | -0.304 | 1.000 |
| **L MFG** | Lexical decision: high vs. low action | -0.508 | 0.585 | -0.867 | 1.000 |
| **L MFG** | Sound judgment: high vs. low sound | 5.977 | 0.569 | 10.498 | **<0.001** |
| **L MFG** | Sound judgment: high vs. low action | 1.797 | 0.581 | 3.092 | **0.013** |
| **L MFG** | Action judgment: high vs. low sound | 0.56 | 0.587 | 0.954 | 1.000 |
| **L MFG** | Action judgment: high vs. low action | 4.193 | 0.666 | 6.296 | **<0.001** |
| **L ATL** | Lexical decision: high vs. low sound | 0.833 | 0.649 | 1.284 | 0.724 |
| **L ATL** | Lexical decision: high vs. low action | -0.638 | 0.6 | -1.064 | 1.000 |
| **L ATL** | Sound judgment: high vs. low sound | 2.969 | 0.635 | 4.674 | **<0.001** |
| **L ATL** | Sound judgment: high vs. low action | -0.13 | 0.685 | -0.19 | 1.000 |
| **L ATL** | Action judgment: high vs. low sound | 0.885 | 0.546 | 1.622 | 0.395 |
| **L ATL** | Action judgment: high vs. low action | 2.591 | 0.75 | 3.455 | **0.005** |
| **L pMTG** | Lexical decision: high vs. low sound | -0.078 | 0.674 | -0.116 | 1.000 |
| **L pMTG** | Lexical decision: high vs. low action | -0.56 | 0.605 | -0.925 | 1.000 |
| **L pMTG** | Sound judgment: high vs. low sound | 3.763 | 0.82 | 4.59 | **<0.001** |
| **L pMTG** | Sound judgment: high vs. low action | 1.484 | 0.55 | 2.698 | **0.036** |
| **L pMTG** | Action judgment: high vs. low sound | 0.977 | 0.663 | 1.473 | 0.521 |
| **L pMTG** | Action judgment: high vs. low action | 1.693 | 0.615 | 2.753 | **0.031** |
| **L pIPL** | Lexical decision: high vs. low sound | 0.286 | 0.56 | 0.511 | 1.000 |
| **L pIPL** | Lexical decision: high vs. low action | -0.234 | 0.64 | -0.366 | 1.000 |
| **L pIPL** | Sound judgment: high vs. low sound | 4.062 | 0.698 | 5.822 | **<0.001** |
| **L pIPL** | Sound judgment: high vs. low action | 1.341 | 0.573 | 2.34 | 0.086 |
| **L pIPL** | Action judgment: high vs. low sound | 1.862 | 0.706 | 2.639 | **0.042** |
| **L pIPL** | Action judgment: high vs. low action | 2.852 | 0.77 | 3.704 | **0.002** |
| **PCC/Precuneus** | Lexical decision: high vs. low sound | -0.195 | 0.638 | -0.306 | 1.000 |
| **PCC/Precuneus** | Lexical decision: high vs. low action | 0.404 | 0.643 | 0.628 | 1.000 |

| | | | | | |
|---|---|---|---|---|---|
| **PCC/Precuneus** | Sound judgment: high vs. low sound | 5.052 | 0.623 | 8.109 | **<0.001** |
| **PCC/Precuneus** | Sound judgment: high vs. low action | 3.203 | 0.722 | 4.437 | **<0.001** |
| **PCC/Precuneus** | Action judgment: high vs. low sound | 1.667 | 0.658 | 2.532 | 0.054 |
| **PCC/Precuneus** | Action judgment: high vs. low action | 5.625 | 0.811 | 6.94 | **<0.001** |
| **dmPFC** | Lexical decision: high vs. low sound | -0.026 | 0.575 | -0.045 | 1.000 |
| **dmPFC** | Lexical decision: high vs. low action | 0.729 | 0.657 | 1.11 | 0.958 |
| **dmPFC** | Sound judgment: high vs. low sound | 5.156 | 0.693 | 7.443 | **<0.001** |
| **dmPFC** | Sound judgment: high vs. low action | 1.484 | 0.75 | 1.978 | 0.192 |
| **dmPFC** | Action judgment: high vs. low sound | 2.109 | 0.581 | 3.628 | **0.003** |
| **dmPFC** | Action judgment: high vs. low action | 5.807 | 0.848 | 6.845 | **<0.001** |

**Table S15.** Paired t-tests comparing the MVPA decoding accuracies between conditions within each ROI.

| ROI | Comparison | Mean | SD | T | p (corr.) |
|---|---|---|---|---|---|
| **L aIFG** | Lexical decision vs. action judgment for high vs. low action | -3.698 | 5.419 | -4.316 | **0.001** |
| **L aIFG** | Lexical decision vs. action judgment for high vs. low sound | -2.24 | 5.234 | -2.706 | 0.070 |
| **L aIFG** | Lexical decision vs. sound judgment for high vs. low action | 0.013 | 6.625 | 0.012 | 1.000 |
| **L aIFG** | Lexical decision vs. sound judgment for high vs. low sound | -4.193 | 6.909 | -3.838 | **0.003** |
| **L aIFG** | Lexical decision: sound vs. action | -0.378 | 5.846 | -0.408 | 1.000 |
| **L aIFG** | Sound judgment vs. action judgment for high vs. low action | -3.711 | 4.985 | -4.708 | **<0.001** |
| **L aIFG** | Sound judgment vs. action judgment for high vs. low sound | 1.953 | 5.571 | 2.217 | 0.227 |
| **L aIFG** | Sound judgment: sound vs. action | 3.828 | 6.13 | 3.949 | **0.002** |
| **L aIFG** | Action judgment: sound vs. action | -1.836 | 5.283 | -2.198 | 0.238 |
| **L MFG** | Lexical decision vs. action judgment for high vs. low action | -4.701 | 4.831 | -6.154 | **<0.001** |
| **L MFG** | Lexical decision vs. action judgment for high vs. low sound | -0.742 | 4.917 | -0.955 | 1.000 |
| **L MFG** | Lexical decision vs. sound judgment for high vs. low action | -2.305 | 4.945 | -2.948 | **0.038** |
| **L MFG** | Lexical decision vs. sound judgment for high vs. low sound | -6.159 | 5.416 | -7.192 | **<0.001** |
| **L MFG** | Lexical decision: sound vs. action | 0.326 | 4.28 | 0.481 | 1.000 |
| **L MFG** | Sound judgment vs. action judgment for high vs. low action | -2.396 | 5.276 | -2.872 | **0.046** |
| **L MFG** | Sound judgment vs. action judgment for high vs. low sound | 5.417 | 5.43 | 6.309 | **<0.001** |
| **L MFG** | Sound judgment: sound vs. action | 4.18 | 5.85 | 4.518 | **<0.001** |
| **L MFG** | Action judgment: sound vs. action | -3.633 | 5.167 | -4.446 | **<0.001** |
| **L ATL** | Lexical decision vs. action judgment for high vs. low action | -3.229 | 5.031 | -4.06 | **0.002** |
| **L ATL** | Lexical decision vs. action judgment for high vs. low sound | -0.052 | 5.806 | -0.057 | 1.000 |
| **L ATL** | Lexical decision vs. sound judgment for high vs. low action | -0.508 | 5.485 | -0.586 | 1.000 |
| **L ATL** | Lexical decision vs. sound judgment for high vs. low sound | -2.135 | 5.833 | -2.315 | 0.182 |
| **L ATL** | Lexical decision: sound vs. action | 1.471 | 5.585 | 1.666 | 0.726 |
| **L ATL** | Sound judgment vs. action judgment for high vs. low action | -2.721 | 6.108 | -2.818 | 0.053 |
| **L ATL** | Sound judgment vs. action judgment for high vs. low sound | 2.083 | 5.475 | 2.406 | 0.147 |
| **L ATL** | Sound judgment: sound vs. action | 3.099 | 4.707 | 4.164 | **0.001** |
| **L ATL** | Action judgment: sound vs. action | -1.706 | 6.403 | -1.685 | 0.700 |
| **L pMTG** | Lexical decision vs. action judgment for high vs. low action | -2.253 | 5.158 | -2.762 | 0.061 |
| **L pMTG** | Lexical decision vs. action judgment for high vs. low sound | -1.055 | 5.248 | -1.271 | 1.000 |
| **L pMTG** | Lexical decision vs. sound judgment for high vs. low action | -2.044 | 5.462 | -2.367 | 0.161 |
| **L pMTG** | Lexical decision vs. sound judgment for high vs. low sound | -3.841 | 6.847 | -3.548 | **0.007** |
| **L pMTG** | Lexical decision: sound vs. action | 0.482 | 5.565 | 0.548 | 1.000 |

| | | | | | |
|---|---|---|---|---|---|
| **L pMTG** | Sound judgment vs. action judgment for high vs. low action | -0.208 | 4.609 | -0.286 | 1.000 |
| **L pMTG** | Sound judgment vs. action judgment for high vs. low sound | 2.786 | 6.215 | 2.836 | 0.050 |
| **L pMTG** | Sound judgment: sound vs. action | 2.279 | 5.892 | 2.446 | 0.133 |
| **L pMTG** | Action judgment: sound vs. action | -0.716 | 6.163 | -0.735 | 1.000 |
| **L pIPL** | Lexical decision vs. action judgment for high vs. low action | -3.086 | 5.874 | -3.323 | **0.014** |
| **L pIPL** | Lexical decision vs. action judgment for high vs. low sound | -1.576 | 5.078 | -1.962 | 0.398 |
| **L pIPL** | Lexical decision vs. sound judgment for high vs. low action | -1.576 | 4.996 | -1.994 | 0.372 |
| **L pIPL** | Lexical decision vs. sound judgment for high vs. low sound | -3.776 | 5.22 | -4.575 | **<0.001** |
| **L pIPL** | Lexical decision: sound vs. action | 0.521 | 3.739 | 0.881 | 1.000 |
| **L pIPL** | Sound judgment vs. action judgment for high vs. low action | -1.51 | 5.375 | -1.777 | 0.583 |
| **L pIPL** | Sound judgment vs. action judgment for high vs. low sound | 2.201 | 6.619 | 2.103 | 0.294 |
| **L pIPL** | Sound judgment: sound vs. action | 2.721 | 5.968 | 2.884 | **0.045** |
| **L pIPL** | Action judgment: sound vs. action | -0.99 | 5.943 | -1.053 | 1.000 |
| **PCC/Precuneus** | Lexical decision vs. action judgment for high vs. low action | -5.221 | 6.492 | -5.087 | **<0.001** |
| **PCC/Precuneus** | Lexical decision vs. action judgment for high vs. low sound | -1.862 | 6.636 | -1.774 | 0.587 |
| **PCC/Precuneus** | Lexical decision vs. sound judgment for high vs. low action | -2.799 | 5.701 | -3.105 | **0.025** |
| **PCC/Precuneus** | Lexical decision vs. sound judgment for high vs. low sound | -5.247 | 5.926 | -5.6 | **<0.001** |
| **PCC/Precuneus** | Lexical decision: sound vs. action | -0.599 | 4.314 | -0.878 | 1.000 |
| **PCC/Precuneus** | Sound judgment vs. action judgment for high vs. low action | -2.422 | 6.684 | -2.292 | 0.192 |
| **PCC/Precuneus** | Sound judgment vs. action judgment for high vs. low sound | 3.385 | 5.149 | 4.158 | **0.001** |
| **PCC/Precuneus** | Sound judgment: sound vs. action | 1.849 | 5.349 | 2.186 | 0.244 |
| **PCC/Precuneus** | Action judgment: sound vs. action | -3.958 | 6.132 | -4.083 | **0.001** |
| **dmPFC** | Lexical decision vs. action judgment for high vs. low action | -5.078 | 6.277 | -5.116 | **<0.001** |
| **dmPFC** | Lexical decision vs. action judgment for high vs. low sound | -2.135 | 5.095 | -2.651 | 0.081 |
| **dmPFC** | Lexical decision vs. sound judgment for high vs. low action | -0.755 | 6.774 | -0.705 | 1.000 |
| **dmPFC** | Lexical decision vs. sound judgment for high vs. low sound | -5.182 | 5.722 | -5.728 | **<0.001** |
| **dmPFC** | Lexical decision: sound vs. action | -0.755 | 5.223 | -0.914 | 1.000 |
| **dmPFC** | Sound judgment vs. action judgment for high vs. low action | -4.323 | 6.645 | -4.114 | **0.001** |
| **dmPFC** | Sound judgment vs. action judgment for high vs. low sound | 3.047 | 5.647 | 3.412 | **0.011** |
| **dmPFC** | Sound judgment: sound vs. action | 3.672 | 6.974 | 3.33 | **0.013** |
| **dmPFC** | Action judgment: sound vs. action | -3.698 | 6.672 | -3.505 | **0.008** |

#### MVPA decoding in large-scale functional networks

The following tables report the results of statistical analyses on the decoding accuracies for MVPA decoding in the large-scale functional networks of Yeo et al. (2011). For statistical inference, we first conducted one-sample t-tests for above-chance decoding in each network. Second, we performed paired t-tests comparing the decoding accuracies for each condition between networks. Finally, we conducted paired t-tests comparing the decoding accuracies between conditions within each network. P-values were corrected for multiple comparisons using Bonferroni correction for the number of networks.

**Table S16.** One-sample t-tests on the decoding accuracies of each network against chance level (50%).

| Network | Analysis | Mean | SD | T | p (corr.) |
|---|---|---|---|---|---|
| Visual | Lexical decision: high vs. low sound | 0.625 | 0.693 | 0.902 | 1.000 |
| Visual | Lexical decision: high vs. low action | -0.195 | 0.733 | -0.266 | 1.000 |
| Visual | Sound judgment: high vs. low sound | 2.76 | 0.617 | 4.473 | **<0.001** |
| Visual | Sound judgment: high vs. low action | 1.901 | 0.583 | 3.26 | **0.008** |
| Visual | Action judgment: high vs. low sound | 0.651 | 0.613 | 1.061 | 1.000 |
| Visual | Action judgment: high vs. low action | 3.49 | 0.664 | 5.254 | **<0.001** |
| Somatomotor | Lexical decision: high vs. low sound | 0.326 | 0.627 | 0.519 | 1.000 |
| Somatomotor | Lexical decision: high vs. low action | 0.638 | 0.632 | 1.01 | 1.000 |
| Somatomotor | Sound judgment: high vs. low sound | 3.281 | 0.68 | 4.824 | **<0.001** |
| Somatomotor | Sound judgment: high vs. low action | -0.443 | 0.626 | -0.707 | 1.000 |
| Somatomotor | Action judgment: high vs. low sound | 1.732 | 0.749 | 2.313 | 0.091 |
| Somatomotor | Action judgment: high vs. low action | 3.672 | 0.755 | 4.865 | **<0.001** |
| Dorsal Attention | Lexical decision: high vs. low sound | 0.703 | 0.599 | 1.174 | 0.866 |
| Dorsal Attention | Lexical decision: high vs. low action | -0.326 | 0.688 | -0.473 | 1.000 |
| Dorsal Attention | Sound judgment: high vs. low sound | 6.445 | 0.783 | 8.232 | **<0.001** |
| Dorsal Attention | Sound judgment: high vs. low action | 3.164 | 0.57 | 5.55 | **<0.001** |
| Dorsal Attention | Action judgment: high vs. low sound | 2.266 | 0.665 | 3.408 | **0.005** |
| Dorsal Attention | Action judgment: high vs. low action | 6.667 | 0.786 | 8.487 | **<0.001** |
| Salience Ventral Attention | Lexical decision: high vs. low sound | 0.911 | 0.673 | 1.355 | 0.642 |
| Salience Ventral Attention | Lexical decision: high vs. low action | 0.104 | 0.737 | 0.141 | 1.000 |
| Salience Ventral Attention | Sound judgment: high vs. low sound | 3.984 | 0.637 | 6.253 | **<0.001** |
| Salience Ventral Attention | Sound judgment: high vs. low action | 0.964 | 0.509 | 1.894 | 0.230 |
| Salience Ventral Attention | Action judgment: high vs. low sound | 2.044 | 0.705 | 2.901 | **0.021** |
| Salience Ventral Attention | Action judgment: high vs. low action | 4.062 | 0.724 | 5.611 | **<0.001** |
| Limbic | Lexical decision: high vs. low sound | 0.781 | 0.543 | 1.439 | 0.554 |
| Limbic | Lexical decision: high vs. low action | -0.911 | 0.704 | -1.294 | 1.000 |
| Limbic | Sound judgment: high vs. low sound | 2.591 | 0.686 | 3.778 | **0.002** |
| Limbic | Sound judgment: high vs. low action | 0.534 | 0.575 | 0.928 | 1.000 |
| Limbic | Action judgment: high vs. low sound | 1.055 | 0.644 | 1.637 | 0.384 |
| Limbic | Action judgment: high vs. low action | 2.266 | 0.557 | 4.071 | **0.001** |
| Frontoparietal Control | Lexical decision: high vs. low sound | 0.417 | 0.561 | 0.743 | 1.000 |
| Frontoparietal Control | Lexical decision: high vs. low action | -0.352 | 0.699 | -0.503 | 1.000 |
| Frontoparietal Control | Sound judgment: high vs. low sound | 7.526 | 0.749 | 10.052 | **<0.001** |
| Frontoparietal Control | Sound judgment: high vs. low action | 2.812 | 0.666 | 4.221 | **<0.001** |
| Frontoparietal Control | Action judgment: high vs. low sound | 2.708 | 0.583 | 4.643 | **<0.001** |
| Frontoparietal Control | Action judgment: high vs. low action | 6.094 | 0.722 | 8.445 | **<0.001** |
| Default | Lexical decision: high vs. low sound | -0.13 | 0.487 | -0.267 | 1.000 |
| Default | Lexical decision: high vs. low action | -0.065 | 0.669 | -0.097 | 1.000 |
| Default | Sound judgment: high vs. low sound | 6.536 | 0.745 | 8.776 | **<0.001** |
| Default | Sound judgment: high vs. low action | 4.036 | 0.674 | 5.988 | **<0.001** |

| | | | | | |
|---|---|---|---|---|---|
| Default | Action judgment: high vs. low sound | 2.135 | 0.63 | 3.387 | **0.006** |
| Default | Action judgment: high vs. low action | 6.276 | 0.797 | 7.876 | **<0.001** |

**Table S17.** Paired t-tests comparing the decoding accuracies for each condition between networks.

| **Analysis** | **Comparison** | **Mean** | **SD** | **T** | **p (corr.)** |
|---|---|---|---|---|---|
| Action judgment: high vs. low action | Control vs. Default | -0.182 | 3.537 | -0.326 | 1.000 |
| Action judgment: high vs. low action | Dorsal Attention vs. Control | 0.573 | 4.01 | 0.904 | 1.000 |
| Action judgment: high vs. low action | Dorsal Attention vs. Default | 0.391 | 3.904 | 0.633 | 1.000 |
| Action judgment: high vs. low action | Dorsal Attention vs. Limbic | 4.401 | 4.3 | 6.474 | **<0.001** |
| Action judgment: high vs. low action | Dorsal vs. Ventral Attention | 2.604 | 4.459 | 3.694 | **0.005** |
| Action judgment: high vs. low action | Limbic vs. Control | -3.828 | 4.144 | -5.842 | **<0.001** |
| Action judgment: high vs. low action | Limbic vs. Default | -4.01 | 4.2 | -6.039 | **<0.001** |
| Action judgment: high vs. low action | Somatomotor vs. Control | -2.422 | 3.766 | -4.067 | **0.001** |
| Action judgment: high vs. low action | Somatomotor vs. Default | -2.604 | 4.298 | -3.832 | **0.009** |
| Action judgment: high vs. low action | Somatomotor vs. Dorsal Attention | -2.995 | 3.95 | -4.795 | **<0.001** |
| Action judgment: high vs. low action | Somatomotor vs. Limbic | 1.406 | 3.382 | 2.63 | 0.281 |
| Action judgment: high vs. low action | Somatomotor vs. Ventral Attention | -0.391 | 3.752 | -0.659 | 1.000 |
| Action judgment: high vs. low action | Ventral Attention vs. Control | -2.031 | 3.843 | -3.343 | **0.022** |
| Action judgment: high vs. low action | Ventral Attention vs. Default | -2.214 | 3.922 | -3.57 | **0.007** |
| Action judgment: high vs. low action | Ventral Attention vs. Limbic | 1.797 | 3.447 | 3.297 | **0.015** |
| Action judgment: high vs. low action | Visual vs. Control | -2.604 | 3.947 | -4.173 | **0.001** |
| Action judgment: high vs. low action | Visual vs. Default | -2.786 | 4.002 | -4.403 | **<0.001** |
| Action judgment: high vs. low action | Visual vs. Dorsal Attention | -3.177 | 2.915 | -6.893 | **<0.001** |
| Action judgment: high vs. low action | Visual vs. Limbic | 1.224 | 3.438 | 2.252 | 0.085 |
| Action judgment: high vs. low action | Visual vs. Somatomotor | -0.182 | 3.923 | -0.294 | 1.000 |
| Action judgment: high vs. low action | Visual vs. Ventral Attention | -0.573 | 4.213 | -0.86 | 1.000 |
| Action judgment: high vs. low sound | Control vs. Default | 0.573 | 3.125 | 1.16 | 1.000 |
| Action judgment: high vs. low sound | Dorsal Attention vs. Control | -0.443 | 3.846 | -0.728 | 1.000 |
| Action judgment: high vs. low sound | Dorsal Attention vs. Default | 0.13 | 3.296 | 0.25 | 1.000 |
| Action judgment: high vs. low sound | Dorsal Attention vs. Limbic | 1.211 | 4.215 | 1.817 | 0.539 |
| Action judgment: high vs. low sound | Dorsal vs. Ventral Attention | 0.221 | 3.843 | 0.364 | 1.000 |
| Action judgment: high vs. low sound | Limbic vs. Control | -1.654 | 4.151 | -2.519 | 0.112 |
| Action judgment: high vs. low sound | Limbic vs. Default | -1.081 | 4.243 | -1.611 | 0.807 |
| Action judgment: high vs. low sound | Somatomotor vs. Control | -0.977 | 3.128 | -1.975 | 0.835 |
| Action judgment: high vs. low sound | Somatomotor vs. Default | -0.404 | 3.697 | -0.691 | 1.000 |
| Action judgment: high vs. low sound | Somatomotor vs. Dorsal Attention | -0.534 | 3.738 | -0.903 | 1.000 |
| Action judgment: high vs. low sound | Somatomotor vs. Limbic | 0.677 | 5.566 | 0.769 | 1.000 |
| Action judgment: high vs. low sound | Somatomotor vs. Ventral Attention | -0.313 | 3.872 | -0.51 | 1.000 |
| Action judgment: high vs. low sound | Ventral Attention vs. Control | -0.664 | 3.887 | -1.081 | 1.000 |
| Action judgment: high vs. low sound | Ventral Attention vs. Default | -0.091 | 3.442 | -0.167 | 1.000 |
| Action judgment: high vs. low sound | Ventral Attention vs. Limbic | 0.99 | 4.725 | 1.325 | 1.000 |
| Action judgment: high vs. low sound | Visual vs. Control | -2.057 | 4.123 | -3.156 | **0.013** |
| Action judgment: high vs. low sound | Visual vs. Default | -1.484 | 4.222 | -2.223 | 0.382 |
| Action judgment: high vs. low sound | Visual vs. Dorsal Attention | -1.615 | 4.343 | -2.351 | 0.224 |
| Action judgment: high vs. low sound | Visual vs. Limbic | -0.404 | 4.928 | -0.518 | 1.000 |
| Action judgment: high vs. low sound | Visual vs. Somatomotor | -1.081 | 4.762 | -1.435 | 0.875 |
| Action judgment: high vs. low sound | Visual vs. Ventral Attention | -1.393 | 4.445 | -1.982 | 0.376 |
| Lexical decision: high vs. low action | Control vs. Default | -0.286 | 3.173 | -0.571 | 1.000 |
| Lexical decision: high vs. low action | Dorsal Attention vs. Control | 0.026 | 4.152 | 0.04 | 1.000 |
| Lexical decision: high vs. low action | Dorsal Attention vs. Default | -0.26 | 4.026 | -0.409 | 1.000 |
| Lexical decision: high vs. low action | Dorsal Attention vs. Limbic | 0.586 | 5.182 | 0.715 | 1.000 |
| Lexical decision: high vs. low action | Dorsal vs. Ventral Attention | -0.43 | 4.667 | -0.582 | 1.000 |
| Lexical decision: high vs. low action | Limbic vs. Control | -0.56 | 4.426 | -0.8 | 1.000 |
| Lexical decision: high vs. low action | Limbic vs. Default | -0.846 | 4.239 | -1.263 | 1.000 |
| Lexical decision: high vs. low action | Somatomotor vs. Control | 0.99 | 3.878 | 1.614 | 1.000 |
| Lexical decision: high vs. low action | Somatomotor vs. Default | 0.703 | 4.121 | 1.079 | 1.000 |
| Lexical decision: high vs. low action | Somatomotor vs. Dorsal Attention | 0.964 | 4.546 | 1.341 | 0.736 |
| Lexical decision: high vs. low action | Somatomotor vs. Limbic | 1.549 | 4.617 | 2.122 | 0.429 |
| Lexical decision: high vs. low action | Somatomotor vs. Ventral Attention | 0.534 | 4.299 | 0.785 | 1.000 |
| Lexical decision: high vs. low action | Ventral Attention vs. Control | 0.456 | 3.923 | 0.735 | 1.000 |
| Lexical decision: high vs. low action | Ventral Attention vs. Default | 0.169 | 4.166 | 0.257 | 1.000 |

| | | | | | |
|---|---|---|---|---|---|
| Lexical decision: high vs. low action | Ventral Attention vs. Limbic | 1.016 | 4.115 | 1.561 | 1.000 |
| Lexical decision: high vs. low action | Visual vs. Control | 0.156 | 4.324 | 0.229 | 1.000 |
| Lexical decision: high vs. low action | Visual vs. Default | -0.13 | 4.288 | -0.192 | 1.000 |
| Lexical decision: high vs. low action | Visual vs. Dorsal Attention | 0.13 | 4.908 | 0.168 | 1.000 |
| Lexical decision: high vs. low action | Visual vs. Limbic | 0.716 | 4.928 | 0.919 | 1.000 |
| Lexical decision: high vs. low action | Visual vs. Somatomotor | -0.833 | 4.812 | -1.095 | 1.000 |
| Lexical decision: high vs. low action | Visual vs. Ventral Attention | -0.299 | 4.698 | -0.403 | 1.000 |
| Lexical decision: high vs. low sound | Control vs. Default | 0.547 | 3.532 | 0.979 | 1.000 |
| Lexical decision: high vs. low sound | Dorsal Attention vs. Control | 0.286 | 3.457 | 0.524 | 1.000 |
| Lexical decision: high vs. low sound | Dorsal Attention vs. Default | 0.833 | 3.552 | 1.484 | 0.807 |
| Lexical decision: high vs. low sound | Dorsal Attention vs. Limbic | -0.078 | 4.43 | -0.112 | 1.000 |
| Lexical decision: high vs. low sound | Dorsal vs. Ventral Attention | -0.208 | 4.343 | -0.303 | 1.000 |
| Lexical decision: high vs. low sound | Limbic vs. Control | 0.365 | 4.608 | 0.5 | 1.000 |
| Lexical decision: high vs. low sound | Limbic vs. Default | 0.911 | 3.474 | 1.659 | 0.886 |
| Lexical decision: high vs. low sound | Somatomotor vs. Control | -0.091 | 4.419 | -0.13 | 1.000 |
| Lexical decision: high vs. low sound | Somatomotor vs. Default | 0.456 | 3.828 | 0.753 | 1.000 |
| Lexical decision: high vs. low sound | Somatomotor vs. Dorsal Attention | -0.378 | 3.802 | -0.628 | 1.000 |
| Lexical decision: high vs. low sound | Somatomotor vs. Limbic | -0.456 | 5.107 | -0.564 | 1.000 |
| Lexical decision: high vs. low sound | Somatomotor vs. Ventral Attention | -0.586 | 4.448 | -0.833 | 1.000 |
| Lexical decision: high vs. low sound | Ventral Attention vs. Control | 0.495 | 4.254 | 0.736 | 1.000 |
| Lexical decision: high vs. low sound | Ventral Attention vs. Default | 1.042 | 4.043 | 1.63 | 0.802 |
| Lexical decision: high vs. low sound | Ventral Attention vs. Limbic | 0.13 | 4.447 | 0.185 | 1.000 |
| Lexical decision: high vs. low sound | Visual vs. Control | 0.208 | 4.601 | 0.286 | 1.000 |
| Lexical decision: high vs. low sound | Visual vs. Default | 0.755 | 3.88 | 1.231 | 1.000 |
| Lexical decision: high vs. low sound | Visual vs. Dorsal Attention | -0.078 | 4.323 | -0.114 | 1.000 |
| Lexical decision: high vs. low sound | Visual vs. Limbic | -0.156 | 4.435 | -0.223 | 1.000 |
| Lexical decision: high vs. low sound | Visual vs. Somatomotor | 0.299 | 4.158 | 0.456 | 1.000 |
| Lexical decision: high vs. low sound | Visual vs. Ventral Attention | -0.286 | 4.404 | -0.411 | 1.000 |
| Sound judgment: high vs. low action | Control vs. Default | -1.224 | 3.891 | -1.989 | 0.154 |
| Sound judgment: high vs. low action | Dorsal Attention vs. Control | 0.352 | 3.476 | 0.64 | 1.000 |
| Sound judgment: high vs. low action | Dorsal Attention vs. Default | -0.872 | 3.519 | -1.568 | 0.388 |
| Sound judgment: high vs. low action | Dorsal Attention vs. Limbic | 2.63 | 3.88 | 4.288 | **0.005** |
| Sound judgment: high vs. low action | Dorsal vs. Ventral Attention | 2.201 | 3.494 | 3.983 | **<0.001** |
| Sound judgment: high vs. low action | Limbic vs. Control | -2.279 | 4.163 | -3.462 | **0.003** |
| Sound judgment: high vs. low action | Limbic vs. Default | -3.503 | 3.912 | -5.663 | **<0.001** |
| Sound judgment: high vs. low action | Somatomotor vs. Control | -3.255 | 3.97 | -5.186 | **<0.001** |
| Sound judgment: high vs. low action | Somatomotor vs. Default | -4.479 | 4.807 | -5.894 | **<0.001** |
| Sound judgment: high vs. low action | Somatomotor vs. Dorsal Attention | -3.607 | 4.172 | -5.467 | **<0.001** |
| Sound judgment: high vs. low action | Somatomotor vs. Limbic | -0.977 | 4.218 | -1.464 | 0.921 |
| Sound judgment: high vs. low action | Somatomotor vs. Ventral Attention | -1.406 | 3.977 | -2.236 | 0.218 |
| Sound judgment: high vs. low action | Ventral Attention vs. Control | -1.849 | 4.084 | -2.863 | **0.047** |
| Sound judgment: high vs. low action | Ventral Attention vs. Default | -3.073 | 4.728 | -4.111 | **0.001** |
| Sound judgment: high vs. low action | Ventral Attention vs. Limbic | 0.43 | 3.737 | 0.727 | 1.000 |
| Sound judgment: high vs. low action | Visual vs. Control | -0.911 | 3.742 | -1.54 | 1.000 |
| Sound judgment: high vs. low action | Visual vs. Default | -2.135 | 3.494 | -3.865 | **0.002** |
| Sound judgment: high vs. low action | Visual vs. Dorsal Attention | -1.263 | 3.348 | -2.386 | 0.167 |
| Sound judgment: high vs. low action | Visual vs. Limbic | 1.367 | 4.486 | 1.927 | 0.210 |
| Sound judgment: high vs. low action | Visual vs. Somatomotor | 2.344 | 4.048 | 3.662 | **0.002** |
| Sound judgment: high vs. low action | Visual vs. Ventral Attention | 0.937 | 3.681 | 1.611 | 0.552 |
| Sound judgment: high vs. low sound | Control vs. Default | 0.99 | 3.468 | 1.804 | 0.779 |
| Sound judgment: high vs. low sound | Dorsal Attention vs. Control | -1.081 | 4.056 | -1.685 | 0.699 |
| Sound judgment: high vs. low sound | Dorsal Attention vs. Default | -0.091 | 4.07 | -0.142 | 1.000 |
| Sound judgment: high vs. low sound | Dorsal Attention vs. Limbic | 3.854 | 4.915 | 4.959 | **<0.001** |
| Sound judgment: high vs. low sound | Dorsal vs. Ventral Attention | 2.461 | 3.475 | 4.479 | **0.001** |
| Sound judgment: high vs. low sound | Limbic vs. Control | -4.935 | 3.429 | -9.103 | **<0.001** |
| Sound judgment: high vs. low sound | Limbic vs. Default | -3.945 | 3.287 | -7.59 | **<0.001** |
| Sound judgment: high vs. low sound | Somatomotor vs. Control | -4.245 | 3.795 | -7.073 | **<0.001** |
| Sound judgment: high vs. low sound | Somatomotor vs. Default | -3.255 | 3.861 | -5.332 | **<0.001** |
| Sound judgment: high vs. low sound | Somatomotor vs. Dorsal Attention | -3.164 | 4.473 | -4.474 | **<0.001** |
| Sound judgment: high vs. low sound | Somatomotor vs. Limbic | 0.69 | 3.819 | 1.143 | 1.000 |
| Sound judgment: high vs. low sound | Somatomotor vs. Ventral Attention | -0.703 | 3.354 | -1.326 | 1.000 |
| Sound judgment: high vs. low sound | Ventral Attention vs. Control | -3.542 | 3.638 | -6.158 | **<0.001** |
| Sound judgment: high vs. low sound | Ventral Attention vs. Default | -2.552 | 3.922 | -4.115 | **0.003** |
| Sound judgment: high vs. low sound | Ventral Attention vs. Limbic | 1.393 | 3.993 | 2.207 | 0.233 |

| | | | | | |
|---|---|---|---|---|---|
| Sound judgment: high vs. low sound | Visual vs. Control | -4.766 | 4.921 | -6.125 | **<0.001** |
| Sound judgment: high vs. low sound | Visual vs. Default | -3.776 | 4.577 | -5.218 | **<0.001** |
| Sound judgment: high vs. low sound | Visual vs. Dorsal Attention | -3.685 | 5.232 | -4.455 | **0.001** |
| Sound judgment: high vs. low sound | Visual vs. Limbic | 0.169 | 4.662 | 0.23 | 1.000 |
| Sound judgment: high vs. low sound | Visual vs. Somatomotor | -0.521 | 4.396 | -0.749 | 1.000 |
| Sound judgment: high vs. low sound | Visual vs. Ventral Attention | -1.224 | 4.861 | -1.592 | 1.000 |

**Table S18.** Paired t-tests comparing the decoding accuracies between conditions within each network.

| Network | Comparison | Mean | SD | T | p (corr.) |
|---|---|---|---|---|---|
| Visual | Lexical decision vs. action judgment for high vs. low action | -3.685 | 6.855 | -3.400 | **0.011** |
| Visual | Lexical decision vs. action judgment for high vs. low sound | -0.026 | 6.083 | -0.027 | 1.000 |
| Visual | Lexical decision vs. sound judgment for high vs. low action | -2.096 | 5.303 | -2.500 | 0.117 |
| Visual | Lexical decision vs. sound judgment for high vs. low sound | -2.135 | 5.576 | -2.422 | 0.141 |
| Visual | Lexical decision: sound vs. action | 0.820 | 5.474 | 0.948 | 1.000 |
| Visual | Sound judgment vs. action judgment for high vs. low action | -1.589 | 5.504 | -1.825 | 0.529 |
| Visual | Sound judgment vs. action judgment for high vs. low sound | 2.109 | 5.902 | 2.260 | 0.206 |
| Visual | Sound judgment: sound vs. action | 0.859 | 6.009 | 0.905 | 1.000 |
| Visual | Action judgment: sound vs. action | -2.839 | 6.036 | -2.974 | **0.035** |
| Somatomotor | Lexical decision vs. action judgment for high vs. low action | -3.034 | 6.559 | -2.925 | **0.040** |
| Somatomotor | Lexical decision vs. action judgment for high vs. low sound | -1.406 | 6.142 | -1.448 | 1.000 |
| Somatomotor | Lexical decision vs. sound judgment for high vs. low action | 1.081 | 5.266 | 1.298 | 1.000 |
| Somatomotor | Lexical decision vs. sound judgment for high vs. low sound | -2.956 | 6.380 | -2.930 | **0.039** |
| Somatomotor | Lexical decision: sound vs. action | -0.313 | 5.683 | -0.348 | 1.000 |
| Somatomotor | Sound judgment vs. action judgment for high vs. low action | -4.115 | 5.508 | -4.725 | **<0.001** |
| Somatomotor | Sound judgment vs. action judgment for high vs. low sound | 1.549 | 6.074 | 1.613 | 0.803 |
| Somatomotor | Sound judgment: sound vs. action | 3.724 | 6.199 | 3.799 | **0.003** |
| Somatomotor | Action judgment: sound vs. action | -1.940 | 6.272 | -1.956 | 0.403 |
| Dorsal Attention | Lexical decision vs. action judgment for high vs. low action | -6.992 | 6.280 | -7.042 | **<0.001** |
| Dorsal Attention | Lexical decision vs. action judgment for high vs. low sound | -1.563 | 5.475 | -1.805 | 0.552 |
| Dorsal Attention | Lexical decision vs. sound judgment for high vs. low action | -3.490 | 4.755 | -4.642 | **<0.001** |
| Dorsal Attention | Lexical decision vs. sound judgment for high vs. low sound | -5.742 | 5.631 | -6.449 | **<0.001** |
| Dorsal Attention | Lexical decision: sound vs. action | 1.029 | 5.604 | 1.161 | 1.000 |
| Dorsal Attention | Sound judgment vs. action judgment for high vs. low action | -3.503 | 5.471 | -4.049 | **0.002** |
| Dorsal Attention | Sound judgment vs. action judgment for high vs. low sound | 4.180 | 5.539 | 4.772 | **<0.001** |
| Dorsal Attention | Sound judgment: sound vs. action | 3.281 | 5.923 | 3.503 | **0.008** |
| Dorsal Attention | Action judgment: sound vs. action | -4.401 | 6.822 | -4.080 | **0.002** |
| Salience Ventral Attention | Lexical decision vs. action judgment for high vs. low action | -3.958 | 6.416 | -3.902 | **0.003** |
| Salience Ventral Attention | Lexical decision vs. action judgment for high vs. low sound | -1.133 | 6.124 | -1.170 | 1.000 |
| Salience Ventral Attention | Lexical decision vs. sound judgment for high vs. low action | -0.859 | 6.056 | -0.897 | 1.000 |
| Salience Ventral Attention | Lexical decision vs. sound judgment for high vs. low sound | -3.073 | 5.669 | -3.429 | **0.010** |
| Salience Ventral Attention | Lexical decision: sound vs. action | 0.807 | 6.073 | 0.841 | 1.000 |
| Salience Ventral Attention | Sound judgment vs. action judgment for high vs. low action | -3.099 | 5.774 | -3.394 | **0.011** |
| Salience Ventral Attention | Sound judgment vs. action judgment for high vs. low sound | 1.940 | 5.756 | 2.132 | 0.276 |
| Salience Ventral Attention | Sound judgment: sound vs. action | 3.021 | 5.145 | 3.713 | **0.004** |

| | | | | | |
|---|---|---|---|---|---|
| Salience Ventral Attention | Action judgment: sound vs. action | -2.018 | 6.714 | -1.901 | 0.453 |
| Limbic | Lexical decision vs. action judgment for high vs. low action | -3.177 | 5.771 | -3.482 | **0.009** |
| Limbic | Lexical decision vs. action judgment for high vs. low sound | -0.273 | 5.482 | -0.315 | 1.000 |
| Limbic | Lexical decision vs. sound judgment for high vs. low action | -1.445 | 6.093 | -1.500 | 0.991 |
| Limbic | Lexical decision vs. sound judgment for high vs. low sound | -1.810 | 5.766 | -1.985 | 0.379 |
| Limbic | Lexical decision: sound vs. action | 1.693 | 4.828 | 2.217 | 0.227 |
| Limbic | Sound judgment vs. action judgment for high vs. low action | -1.732 | 3.729 | -2.937 | **0.039** |
| Limbic | Sound judgment vs. action judgment for high vs. low sound | 1.536 | 5.865 | 1.657 | 0.739 |
| Limbic | Sound judgment: sound vs. action | 2.057 | 5.018 | 2.593 | 0.093 |
| Limbic | Action judgment: sound vs. action | -1.211 | 5.299 | -1.445 | 1.000 |
| Frontoparietal Control | Lexical decision vs. action judgment for high vs. low action | -6.445 | 5.597 | -7.283 | **<0.001** |
| Frontoparietal Control | Lexical decision vs. action judgment for high vs. low sound | -2.292 | 5.138 | -2.821 | 0.052 |
| Frontoparietal Control | Lexical decision vs. sound judgment for high vs. low action | -3.164 | 5.860 | -3.415 | **0.011** |
| Frontoparietal Control | Lexical decision vs. sound judgment for high vs. low sound | -7.109 | 5.833 | -7.709 | **<0.001** |
| Frontoparietal Control | Lexical decision: sound vs. action | 0.768 | 5.776 | 0.841 | 1.000 |
| Frontoparietal Control | Sound judgment vs. action judgment for high vs. low action | -3.281 | 6.022 | -3.446 | **0.010** |
| Frontoparietal Control | Sound judgment vs. action judgment for high vs. low sound | 4.818 | 5.594 | 5.447 | **<0.001** |
| Frontoparietal Control | Sound judgment: sound vs. action | 4.714 | 6.500 | 4.586 | **<0.001** |
| Frontoparietal Control | Action judgment: sound vs. action | -3.385 | 5.546 | -3.861 | 0.003 |
| Default | Lexical decision vs. action judgment for high vs. low action | -6.341 | 6.358 | -6.308 | **<0.001** |
| Default | Lexical decision vs. action judgment for high vs. low sound | -2.266 | 5.196 | -2.758 | 0.062 |
| Default | Lexical decision vs. sound judgment for high vs. low action | -4.102 | 5.254 | -4.937 | **<0.001** |
| Default | Lexical decision vs. sound judgment for high vs. low sound | -6.667 | 5.786 | -7.288 | **<0.001** |
| Default | Lexical decision: sound vs. action | -0.065 | 4.962 | -0.083 | 1.000 |
| Default | Sound judgment vs. action judgment for high vs. low action | -2.240 | 6.201 | -2.284 | 0.195 |
| Default | Sound judgment vs. action judgment for high vs. low sound | 4.401 | 5.671 | 4.908 | **<0.001** |
| Default | Sound judgment: sound vs. action | 2.500 | 6.586 | 2.401 | 0.149 |
| Default | Action judgment: sound vs. action | -4.141 | 5.830 | -4.492 | **<0.001** |

#### Cross-decoding of task-relevant conceptual features

We also performed cross-decoding in each network of the 7-network parcellation by Yeo et al. (2011), training on the activity patterns for [sound judgment: high vs. low sound words], and testing on the activity patterns for [action judgment: high vs. low action words], as well as vice versa.

The following tables report the results of statistical analyses on the decoding accuracies. We first conducted one-sample t-tests for above-chance decoding in each network. Second, we performed paired t-tests comparing the decoding accuracies between networks. P-values were corrected for multiple comparisons using Bonferroni correction for the number of networks.

**Table S19.** One-sample t-tests on the decoding accuracies of each network against chance level (50%).

| Network | Mean | SD | T | p (corr.) |
|---|---|---|---|---|
| Visual | 1.133 | 0.361 | 3.139 | **0.011** |
| Somatomotor | 1.745 | 0.426 | 4.099 | **0.001** |
| Dorsal Attention | 2.637 | 0.447 | 5.905 | **<0.001** |
| Salience Ventral Attention | 2.057 | 0.480 | 4.285 | **<0.001** |
| Limbic | 1.016 | 0.487 | 2.084 | 0.153 |
| Frontoparietal Control | 3.828 | 0.521 | 7.351 | **<0.001** |
| Default | 2.826 | 0.547 | 5.164 | **<0.001** |

**Table S20.** Paired t-tests comparing the decoding accuracies between networks.

| Network | Mean | SD | T | p (corr.) |
|---|---|---|---|---|
| Control vs. Default | 1.003 | 2.186 | 2.901 | **0.043** |
| Dorsal Attention vs. Control | -1.191 | 3.383 | -2.227 | 0.222 |
| Dorsal Attention vs. Default | -0.189 | 3.482 | -0.343 | 1.000 |
| Dorsal Attention vs. Limbic | 1.621 | 3.701 | 2.770 | 0.060 |
| Dorsal Attention vs. Ventral Attention | 0.579 | 3.233 | 1.133 | 1.000 |
| Limbic vs. Control | -2.813 | 3.320 | -5.358 | **<0.001** |
| Limbic vs. Default | -1.810 | 3.135 | -3.651 | **0.005** |
| Ventral Attention vs. Control | -1.771 | 2.866 | -3.908 | **0.003** |
| Ventral Attention vs. Default | -0.768 | 2.968 | -1.637 | 0.768 |
| Ventral Attention vs. Limbic | 1.042 | 2.803 | 2.350 | 0.167 |
| Somatomotor vs. Control | -2.083 | 2.987 | -4.412 | **0.001** |
| Somatomotor vs. Default | -1.081 | 3.210 | -2.129 | 0.277 |
| Somatomotor vs. Dorsal Attention | -0.892 | 2.917 | -1.934 | 0.423 |
| Somatomotor vs. Limbic | 0.729 | 3.483 | 1.324 | 1.000 |
| Somatomotor vs. Ventral Attention | -0.312 | 3.129 | -0.632 | 1.000 |
| Visual vs. Control | -2.695 | 2.741 | -6.219 | **<0.001** |
| Visual vs. Default | -1.693 | 3.173 | -3.374 | **0.012** |
| Visual vs. Dorsal Attention | -1.504 | 2.539 | -3.746 | **0.004** |
| Visual vs. Limbic | 0.117 | 3.221 | 0.230 | 1.000 |
| Visual vs. Ventral Attention | -0.924 | 2.612 | -2.238 | 0.217 |
| Visual vs. Somatomotor | -0.612 | 2.525 | -1.533 | 0.933 |
